# LptM promotes oxidative maturation of the lipopolysaccharide translocon by substrate binding mimicry

**DOI:** 10.1101/2023.01.02.522452

**Authors:** Yiying Yang, Haoxiang Chen, Robin A. Corey, Violette Morales, Yves Quentin, Carine Froment, Anne Caumont-Sarcos, Cécile Albenne, Odile Burlet-Schiltz, David Ranava, Phillip J. Stansfeld, Julien Marcoux, Raffaele Ieva

**Affiliations:** Laboratoire de Microbiologie et Génétique Moléculaires, Centre de Biologie Intégrative (CBI), Université de Toulouse, CNRS, UPS, Toulouse 31062, France; Department of Biochemistry, University of Oxford, Oxford, UK; Institut de Pharmacologie et de Biologie Structurale (IPBS), Université de Toulouse, CNRS, UPS, Toulouse 31062, France

## Abstract

Insertion of lipopolysaccharide (LPS) into the outer membrane (OM) of Gram-negative bacteria is mediated by a druggable OM translocon consisting of a β-barrel membrane protein, LptD, and a lipoprotein, LptE. The β-barrel assembly machinery (BAM) assembles LptD together with LptE to form a plug-and-barrel structure. In the enterobacterium *Escherichia coli*, formation of two native disulfide bonds in LptD controls LPS translocon activation. Here we report the discovery of LptM (formerly YifL), a conserved lipoprotein that assembles together with LptD and LptE at the BAM complex. We demonstrate that LptM stabilizes a conformation of LptD that can efficiently acquire native disulfide bonds and be released as mature LPS translocon by the BAM complex. Inactivation of LptM causes the accumulation of non-natively oxidized LptD, making disulfide bond isomerization by DsbC become essential for viability. Our structural prediction and biochemical analyses indicate that LptM binds to sites in both LptD and LptE that are proposed to coordinate LPS insertion into the OM. These results suggest that LptM facilitates oxidative maturation of LptD by mimicking LPS binding, thereby activating the LPS translocon.

## INTRODUCTION

Gram-negative bacteria surround and protect their cytoplasm with a multilayered envelope formed by an inner membrane (IM), an outer membrane (OM) and a thin peptidoglycan layer, sandwiched in between. Both lipid bilayers carry out crucial functions, including nutrient uptake and energy metabolism, as well as cytoplasm detoxification and protection against noxious chemicals, thereby promoting adaptability to a wide range of niches (Nikaido, 2003). The OM forms the foremost barrier to cellular access and therefore, in the context of bacterial multidrug resistance, the OM is a particularly attractive drug target for the development of new antimicrobials (Robinson, 2019; Sousa, 2019).

Lipopolysaccharide (LPS) is a major structural component that accumulates in the external leaflet of the OM lipid bilayer. LPS typically contains six saturated acyl chains, surface displayed carbohydrates and negatively charged phosphate groups that are stabilized by divalent cations. This therefore structures a densely-packed layer that shields the cell from lipophilic molecules, detergents and antimicrobial compounds. LPS is also a modulator of protein homeostasis and peptidoglycan remodeling, regulating the overall architecture of the bacterial envelope (Klein and Raina, 2019; Lima et al., 2013; Morè et al., 2019; Sperandeo et al., 2008). In addition to its protective function, LPS acts as an endotoxin with potent immunogenic activity and represents a major target of the innate immune response (Bertani and Ruiz, 2018).

Anterograde LPS transfer across the bacterial envelope is mediated by a dedicated LPS transport (Lpt) pathway. In the enterobacterial model organism *Escherichia coli*, the Lpt pathway relies on the activity of 7 proteins, known as LptA-G (Okuda et al., 2016; Sperandeo et al., 2019). Impairment of the Lpt pathway causes accumulation of LPS in the IM (Sperandeo et al., 2008), where it undergoes subsequent modification by colanic acid (Meredith et al., 2007), a polysaccharide produced upon envelope damage (Wall et al., 2018).

The terminal link of the Lpt pathway consists of an OM translocon that inserts LPS into the external leaflet of the OM. The OM LPS translocon, composed by an integral OM protein, LptD, and a cognate OM-associated lipoprotein, LptE, acquires a peculiar “plug-and-barrel” architecture (Dong et al., 2014; Freinkman et al., 2011; Qiao et al., 2014). LptD consists of an amino (N)-terminal periplasmic β-taco domain and a carboxy (C)-terminal 26-stranded β-barrel transmembrane domain. The β-taco domain receives LPS from a structurally analogous periplasmic protein, LptA. The C-terminal β-barrel domain defines a large internal lumen that is partly plugged by LptE (Botos et al., 2016; Dong et al., 2014; Qiao et al., 2014). After docking onto the LptD β-taco domain, LPS enters the outer membrane via a mechanism involving a membrane-embedded interface between the β-taco and the β-barrel domains of LptD. This region is proximal to the β-barrel lateral gate that forms between the first and the last β-strands of the LptD transmembrane domain (Botte et al., 2022; Dong et al., 2014; Fiorentino et al., 2021; Gu et al., 2015).

Biogenesis of the OM LPS translocon requires the cooperation of the β-barrel assembly machinery (BAM) and the periplasmic disulfide bond formation machinery (Dsb). BAM is a heteropentamer (BamABCDE) that folds integral OM proteins, such as LptD, into membrane-spanning β-barrel structures (Ranava et al., 2018; Ricci and Silhavy, 2019). The catalytic, integral OM protein BamA and the lipoprotein BamD are two essential BAM subunits that directly coordinate LptD and LptE assembly to form a plug-and-barrel structure (Lee et al., 2016b). Chaperones and proteases help preventing the accumulation of off-pathway folding intermediates that would hamper OMP biogenesis if blocked at the BAM complex (Narita et al., 2013; Soltes et al., 2017). Within LptD, two native disulfide bonds between pairs of non-consecutive cysteine residues interlink the β-taco and the β-barrel domains, thereby activating the LPS translocon (Ruiz et al., 2010). This complex LptD oxidation pattern occurs via a stepwise maturation process. The first two consecutive cysteines in the β-taco domain are initially oxidized by the oxidase DsbA, generating a non-native disulfide bond. After engaging with the BAM complex for assembly of LptD together with LptE in the OM, acquisition of native disulfide bonds in LptD activates the translocon (Chng et al., 2012). Typically, DsbC promotes disulfide bond shuffling in secretory proteins (Gleiter and Bardwell, 2008), however the role of this enzyme in LptD oxidative folding has been debated (Denoncin et al., 2010; Ruiz et al., 2010).

The development of Lpt-targeted drugs requires detailed understanding of the process of LPS transport to the OM. Here we report the discovery of LptM as a novel component of the OM LPS translocon. LptM binds the membrane-embedded portion of the translocon, making contacts with sites in both LptD and LptE that are proposed to mediate LPS insertion into the OM. Thus, LptM stabilizes an active conformation of the OM LPS translocon promoting its assembly by the BAM complex and subsequent oxidative maturation. LptM represent a new type of component in the LPS transport pathway, which controls activation of the OM LPS translocon by mimicking substrate binding.

## RESULTS

### The OM LPS translocon stably interacts with LptM

We discovered that the *E. coli* OM LPS translocon, LptDE, interacts with an uncharacterized lipoprotein, formerly known as YifL, that we have renamed LptM (<u>Lpt</u>D oxidative <u>m</u>aturation-associated lipoprotein). We identified LptM upon expression and Ni-affinity purification of the LptDE^His^ complex (harboring a C-terminally poly-histidine tagged variant of LptE) using a mild, non-ionic detergent. Sodium dodecyl sulfate (SDS)-polyacrylamide gel electrophoresis (PAGE) of the elution fraction followed by Coomassie brilliant blue staining showed efficient co-purification of LptD with the bait protein LptE^His^ (Figure 1A, lane 1). Analysis of the same elution fraction by blue native (BN)-PAGE revealed that the OM LPS translocon migrated with an apparent molecular weight of approximately 140 kDa (Figure 1A, lane 3). To verify the protein content of this major gel band, we employed MALDI-TOF/TOF tandem mass spectrometry (MS) after trypsin digestion. Surprisingly, in addition to LptD and LptE, this analysis identified the uncharacterized lipoprotein LptM (Figure S1). To explore the role of this factor, we purified the LptDE complex from a strain lacking the *yifL/lptM* gene. Although the yields of LptE^His^ and LptD resolved by SDS-PAGE were unchanged (Figure 1A, lanes 1 and 2), the intensity of the blue native 140 kDa band was reduced (Figure 1A, lanes 3 and 4). Part of purified LptE^His^ and LptD originated a slightly slower-migrating BN gel band. Being LptM relatively small in mass (6 kDa) its presence is unlikely to modify the migration of the LptDE complex on BN-gels. The accumulation of a slower-migrating form of the LptDE complex could be indicative of an effect of LptM on the native organization of the OM LPS translocon.

**Figure 1.**
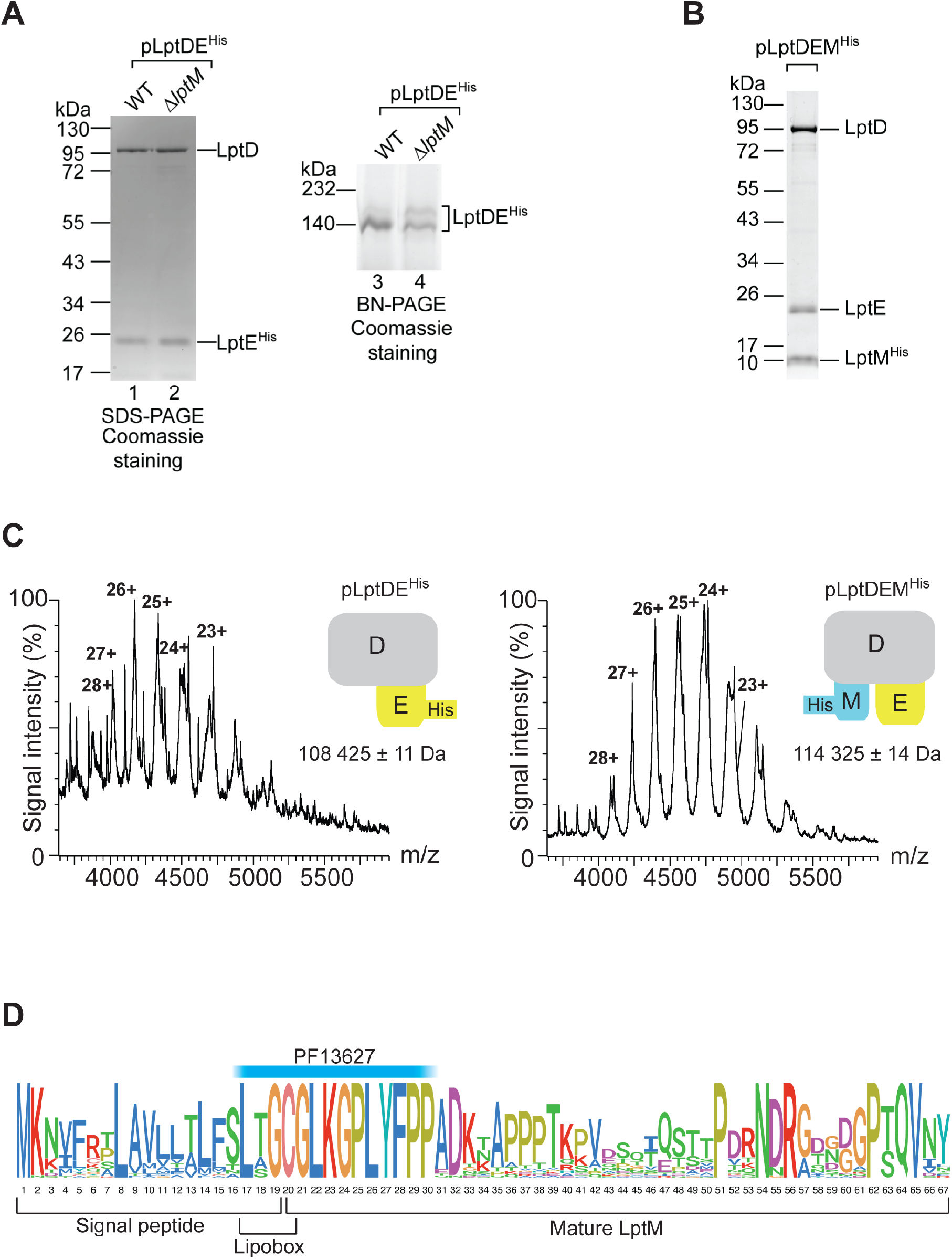
LptM interacts with the LPS translocon. **(A)** The envelope fractions of wild-type (WT, *lptM*^+^) and Δ*lptM* transformed with pLptDE^His^ cells were solubilized using the mild detergent DDM and subjected to Ni-affinity purification. The purified LPS translocon was analyzed by SDS-PAGE (lanes 1 and 2) and BN-PAGE (lanes 3 and 4) followed by Coomassie Brilliant Blue staining. **(B)** Mild detergent-solubilized LptM^His^ purified from Δ*lptM* cells transformed pLptDEM^His^ was analyzed by SDS-PAGE and Coomassie Brilliant Blue staining. **(C)** Native mass-spectrometry analysis of LptDE^His^ and of LptDEM^His^ purified as in (A) and (B), respectively. A schematic diagram of each complex is shown for each spectrum; LptD in grey, LptE in yellow, LptM in light blue. Left: LptDE^His^ formed a heterodimeric complex of 108425 ±11 Da (LptD, theoretical mass of the mature protein = 87080 Da; LptE^His^, theoretical mass of the mature and tri-acylated protein = 21348 Da). Right: LptDEM^His^ formed a heterotrimeric complex of 114325 ± 14 Da (LptD, theoretical mass of the mature protein = 87080 Da; LptE, theoretical mass of the mature and triacylated protein = 20248 Da; LptM^His^, theoretical mass of the mature and triacylated protein = 6988 Da). The mass difference between the two complexes is 5900 Da ± 25 Da, corresponding to the mass of mature tri-acylated LptM 5888 Da. **(D)** Logoplot representation of the LptM amino acid sequence in *Enterobacteriaceae*. The plot was obtained from the multiple alignment of 37 amino acid sequences of LptM in representative genomes of a restricted Enterobacriaceae family reported in Figure S3 (see also Supplementary Information).

To address whether LptM can stably interact with the OM LPS translocon, a C-terminally poly-histidine tagged version of *E. coli* LptM (LptM^His^) was overproduced together with LptD and LptE. Ni-affinity chromatography revealed efficient purification of the OM LPS translocon subunits along with the bait protein (Figure 1B). Most notably, native MS demonstrated that LptD, LptE and LptM^His^ formed a complex which is larger by 5900 Da ± 25 Da than the LptDE^His^ heterodimer (Figure 1C). This mass difference fits well with the theoretical molecular weight of mature tri-acylated LptM, 5888 Da. Together, these results indicate that LptM stably associates with LptDE forming a 1:1:1 heterotrimeric complex.

### LptM is a lipoprotein conserved in *Enterobacteriaceae*

The gene *lptM* codes for a small lipoprotein of approximately 6 kDa. The LptM lipobox cysteine residue is followed by a glycine residue (Figure 1D), suggesting that the protein is transported to the OM (Yamaguchi et al., 1988), in agreement with our finding that LptM interacts with the OM complex LptDE. LptM includes the annotated Pfam domain, PF13627 (Prokaryotic lipoprotein-attachment site), overlapping with the protein lipobox and a short N-terminal region of the mature protein. This motif was used to search *lptM* putative orthologs in a group of 2927 bacterial genomes (including at least one genome per bacterial family of those present in the bacterial Genome Taxonomy Database, GTDB) (Parks et al., 2018). Our results indicate that *lptM*-like genes are present in approximately 79% of Gamma- and 57% of Alpha-proteobacteria (Figure S2, see also Supplementary Information), and is virtually absent in non-proteobacterial genomes. The degree of conservation of chromosomal neighboring genes surrounding *lptM* in proteobacteria suggests inheritance of this gene from a common ancestor. Thus, the identified *lptM* genes can be considered orthologs (Figure S2). The amino acid sequence length downstream PF13627 is generally short with less than 70 amino acids (Figure S2). Both the N-terminal PF13627 and the C-terminal moiety of the protein are conserved in a recently redefined *Enterobacteriaceae* family, whereas other proteobacteria harbor a lipoprotein containing the N-terminal PF13627 motif followed by a C-terminal region that is poorly conserved with that of *E. coli* LptM (Figure 1D and Figure S3; see also Supplementary Information). Taken together, our taxonomic analysis strongly suggests that distinct sub-regions of LptM from *Enterobacteriaceae* are functionally important.

### LptM promotes efficient assembly of the OM LPS translocon at the BAM complex

The biogenesis of the LptDE complex occurs stepwise. The large β-barrel domain of LptD is folded by the BAM complex embedding LptE in its internal lumen. Given our observation that LptM has a strong affinity for the OM LPS translocon, we assessed whether LptM would be required for efficient assembly of LptD together with LptE. To test this hypothesis, first we generated a double deletion strain lacking *lptM* and *bamB*, which encodes a subunit of the BAM complex that is particularly critical for LPS translocon biogenesis (Ruiz et al., 2005). *DbamB* is sensitive to detergents, such as SDS, and to large antibiotics, such as vancomycin, which are normally excluded from the OM of wild-type cells. To compare the phenotypes of single and double *lptM* and *bamB* deletion strains, we used low concentrations of vancomycin and SDS that have a partial inhibitory effect on *DbamB* and that have no effect on Δ*lptM* (Figure 2A, panels 3-6). The sensitivity of the double deletion strain to both SDS and vancomycin was considerably accentuated compared to *DbamB*, especially at the higher temperature of 42°C (Figure 2A, panels 4 and 6). This detrimental synergistic effect reveals a genetic association of *lptM* with *bamB*, pointing to a possible role of LptM in LptDE assembly.

**Figure 2.**
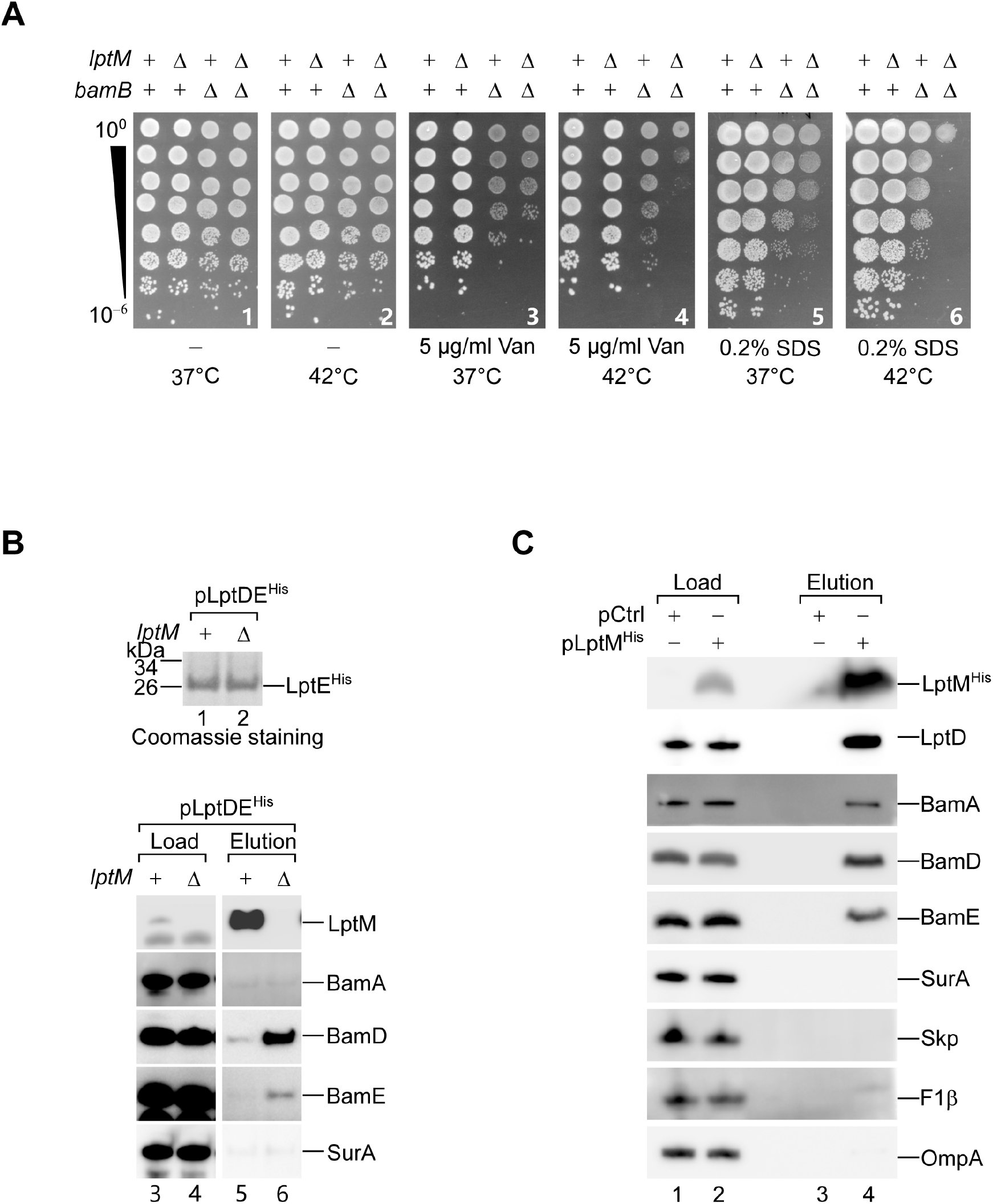
LptM facilitates assembly of the LPS translocon by the BAM complex. **(A)** Drop dilution growth test of *E. coli* wild-type and the indicated derivative strains deleted of *bamB, lptM* or both under the indicated conditions. **(B)** The DDM-solubilized and purified LPS translocon obtained from *lptM^+^* or Δ*lptM* cells harboring pLptDE^His^ was analyzed by Coomassie Brilliant Blue staining to visualize the bait protein (LptE^His^, lanes 1 and 2) and Western blotting using the indicated antisera to determine co-eluted proteins (lanes 3-6). Load: 1.8%; Elution: 100%. **(C)** Mild detergent-solubilized LptM^His^ purified from Δ*lptM* cells transformed with an empty vector pCtrl or pLptM^His^ was analyzed by SDS-PAGE and Western blotting using the indicated antisera. Load: 2%; Elution: 100%.

Mutations that impair the folding of LptD delay LPS translocon assembly into the OM, thereby prolonging its time of residence at the BAM complex (Lee et al., 2019; Lee et al., 2016b). We reasoned that, if LptM was indeed important for efficient assembly of LptDE, the deletion of *lptM* could cause accumulation of the LPS translocon at the BAM complex. To address this hypothesis, LptDE^His^ was produced either in wild-type (*lptM^+^*) or in Δ*lptM* cells and affinity purified, yielding similar amounts of the bait protein (Figure 2B, lanes 1 and 2). Western blot analysis of the elution fractions revealed that LptM was highly enriched in the LptDE^His^ elution fraction obtained from *lptM^+^* cells (Figure 2B, lane 5). Strikingly, the BAM subunits BamD and BamE were co-purified along with LptE^His^ in Δ*lptM* but not in *lptM^+^* cells, indicating a prolonged interaction of LptDE^His^ with the BAM complex (Figure 2B, lanes 5 and 6). We conclude that LptM is required for an optimal step of LPS translocon assembly that occurs prior to translocon release into the OM by the BAM complex. Furthermore, affinity purification of overproduced LptM^His^ copurified LptD and, albeit to a lower extent, also subunits of the BAM complex (Figure 2C, lane 4). This result is consistent with a mechanistic model where LptM associates with LptD and LptE during an early step of their assembly as a plug-and-barrel complex.

Next, we employed site-directed photocrosslinking to probe the environment surrounding LptM in an attempt to map its interactions with the OM LPS translocon. Using an amber suppression method (Chin et al., 2002), the photoactivatable amino acid analog para-benzoyl-phenylalanine (pBpa) was introduced at 6 different positions of the LptM^His^ sequence. Aliquots of cells that expressed the photocrosslinkable LptM variants were split in two halves, one of which was subjected to UV irradiation to activate pBpa. Upon affinity purification and western-blotting with LptM-specific antibodies, a number of UV-dependent crosslink adducts were detected (Figure S4A). The incorporation of pBpa at position L22 and, to a lower extent, Y27 led to the formation of a UV-dependent adduct of ~100 kDa that could potentially correspond to an interaction of LptM with the ~88 kDa LptD protein. LptD contains disulfide bonds that influence its migration on polyacrylamide gels. Thus, probing for the presence of LptD, we found that the SDS-PAGE migration of the L22pBpa crosslink adduct was sensitive to a reducing agent (Figure S4B), indicating the presence of LptD. In addition, two major crosslink adducts of an apparent molecular weight of ~50 kDa were obtained with pBpa introduced in the C-terminal half of LptM (Figure S4A). Based on their apparent mass, we surmised that these crosslinked adducts could correspond to an interaction of LptM with the abundant outer membrane protein OmpA (~37 kDa). This hypothesis was proven correct by conducting photocrosslinking in a *DompA* strain and by using OmpA-specific antibodies (Figure S4C). As OmpA was not detected upon native purification of LptM^His^ (Figure 2C), the observed LptM-OmpA crosslink adducts are presumably due to spurious association of the overproduced bait with the abundant OM protein OmpA. Taken together, our data indicate that the N-terminal region of LptM can directly contact LptD.

### Inactivation of *lptM* impairs LptD oxidative maturation

We asked whether LptM influences the oxidative maturation of LptD that occurs concomitantly to LPS translocon assembly by the BAM complex. LptD contains four cysteine residues at position 31, 173, 724 and 725 of its amino acid sequence, hereafter named C1 to C4, respectively (Figure 3A). Upon transport into the periplasm by the Sec machinery, the two most N-terminal cysteine residues C1 and C2 are oxidized by DsbA, leading to formation of an oxidation intermediate (LptD^C1-C2^). Folding of LptD^C1-C2^ into a β-barrel structure that surrounds LptE (Freinkman et al., 2011; Lee et al., 2016b) is followed by the isomerization of the C1-C2 disulfide bond to the C2-C4 bond and further oxidation of the remaining C1 and C3 cysteines (Chng et al., 2012; Ruiz et al., 2010). Hereafter, we will refer to the mature, fully oxidized form of LptD (LptD^C1-C3,C2-C4^) as LptD^Ox^. At least one disulfide bond between non-consecutive cysteines (C1-C3 or C2-C4) is required for proper LPS translocon function, whereas the earlier assembly intermediate, LptD^C1-C2^, is functionally defective (Chng et al., 2012; Denoncin et al., 2010; Ruiz et al., 2010). Despite being critical for LptD maturation, the mechanisms mediating LptD disulfide bond formation and isomerization remain only partially understood (Denoncin et al., 2010; Meehan et al., 2017; Ruiz et al., 2010).

**Figure 3.**
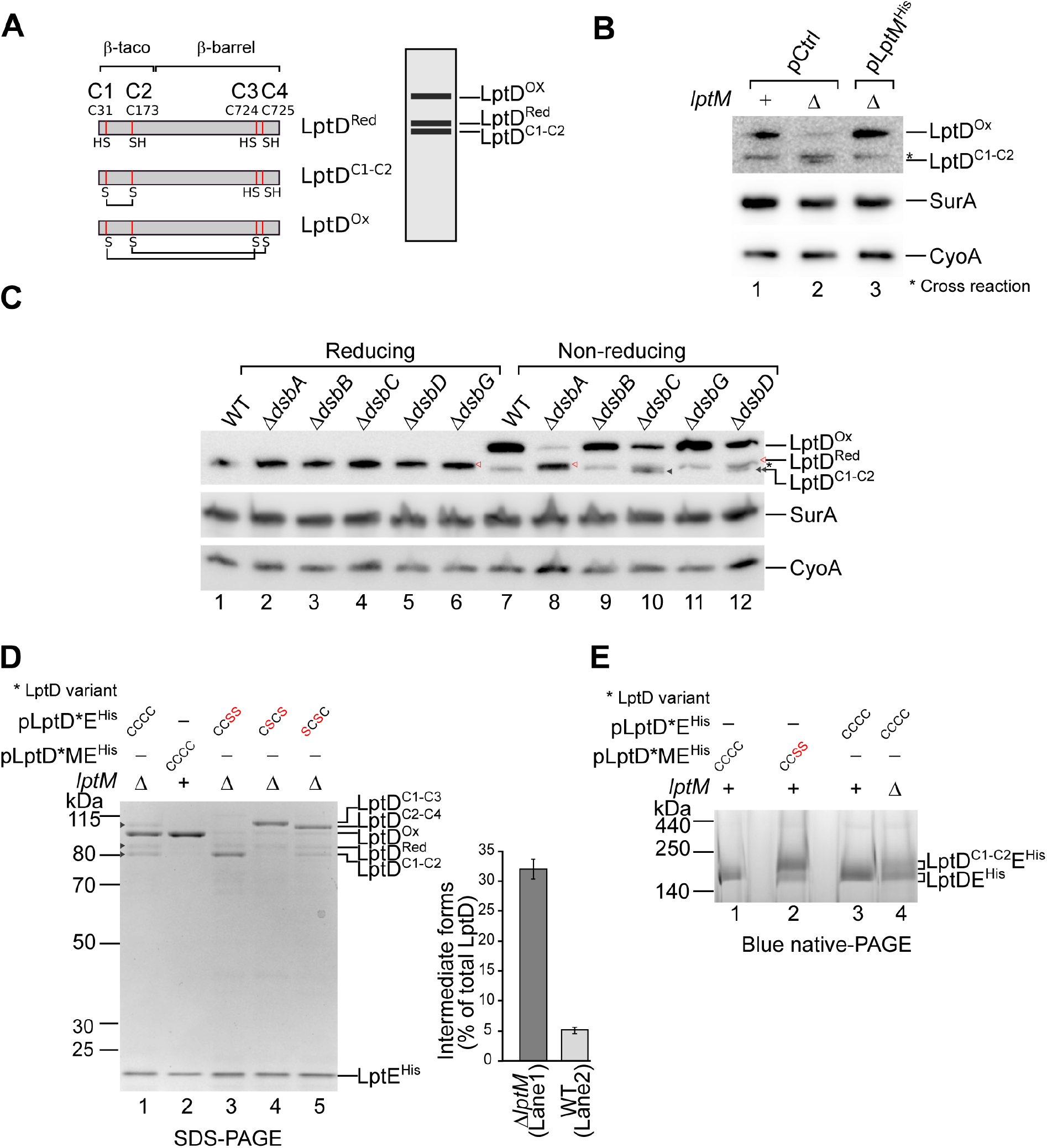
LptM promotes LptD oxidative maturation. **(A)** Schematic representation of mature LptD primary sequence and Cys positions of LptD^Red^ (top, left), as well as the disulfide bonds that characterize LptD^C1-C2^ (middle, left) or fully oxidized LptD^Ox^ (bottom, left). A typical migration pattern corresponding to different oxidative states of LptD by nonreducing SDS-PAGE is schematically represented (right). **(B)** The total protein contents of wild-type (*lptM^+^*) and Δ*lptM* strains transformed with the empty vector pCtrl, and the complementation strain Δ*lptM* transformed with pLptM^His^ were separated by SDS-PAGE and analyzed by Western blotting using the indicated antisera. **(C)** The total protein contents of wild-type and strains lacking single *dsb* genes were analyzed as in (B). **(D)** Purification of LptE^His^ from Δ*lptM* cells that overproduce LptDE^His^ (transformed with pLptDE^His^, lane 1) or from wild-type cells that overproduce LptDE^His^ and LptM (transformed with pLptDME^His^, lane 2) or Δ*lptM* cells that overproduce the indicated Cys-to-Ser mutant forms of LptDE^His^ (transformed with pLptD^CCSS^E^His^ or pLptD^CSCS^E^His^ or pLptD^SCSC^E^His^, respectively lanes 3-5). The signals of any LptD forms in lanes 1 and 2 were quantified. The amount of intermediate forms (LptD^Red^ + LptD^C1-C2^ + LptD^C2-C4^) was normalized to the overall amount of all LptD forms (LptD^Red^ + LptD^C1-C2^ + LptD^C2-C4^ + LptD^ox^). Data are represented as means ± SEM (N = 3). The “ * “ indicates the plasmid-encoded LptD variant, wild-type LptD^CCCC^, LptD^CCSS^, LptD^CSCS^ or LptD^SCSC^, as specified on the top of each gel lane. **(E)** LptDE^His^ containing either wild-type (LptD^CCCC^) or LptD^CCSS^ cooverproduced together with LptM or alone in wild-type or Δ*lptM* cells were Ni-affinity purified and resolved by BN-PAGE, prior to Coomassie Brilliant Blue staining. The “ * “ indicates the plasmid-encoded LptD variant, wild-type LptD^CCCC^, LptD^CCSS^, as specified on the top of each gel lane.

The oxidation states of LptD can be determined by non-reducing SDS-PAGE, as LptD^C1-C2^ migrates slightly faster than reduced LptD (LptD^Red^), whereas LptD^Ox^ migrates slower than both LptD^C1-C2^ and LptD^Red^ (Figure 3A)(Chng et al., 2012; Ruiz et al., 2010). Strikingly, our analysis of endogenous LptD in wild-type and Δ*lptM* cells revealed that the level of LptD^Ox^ was drastically reduced in Δ*lptM* and fully restored in a derivative complementation strain harboring plasmid-encoded LptM^His^ (Figure 3B). Of note, Δ*lptM* cells accumulated some low amount of a faster-migrating form of LptD. To identify this LptD form, we analyzed the migration of LptD produced by strains lacking single components of the periplasmic disulfide bond formation (Dsb) oxidative folding machinery. The lack of the oxidase DsbA caused the accumulation of the reduced form, LptD^Red^ (Figure 3C, lane 8) (Ruiz et al., 2010), whereas elimination of the thiol-quinone oxidoreductase DsbB and the reductase DsbG had no obvious effect on LptD oxidation under these condtions (Figure 3C, lanes 9 and 11). In contrast, the lack of DsbC and of its electron donor DsbD caused the accumulation of LptD^C1-C2^ (Figure 3C, lanes 10 and 12). The LptD migration pattern obtained with the deletion of *lptM* is similar to that obtained with Δ*dsbC* and *DdsbD*, although in Δ*lptM* the amount of LptD^Ox^ was particularly low (Figure 3B, lane 2). Taken together, these results indicate that a Δ*lptM* strain fails to efficiently form mature LptD^Ox^ and accumulates the assembly intermediate, LptD^C1-C2^.

To further analyze the LPS translocon maturation defect caused by the absence of LptM, we overproduced LptD and LptE^His^ either with or without LptM and performed Ni-affinity purifications. As a reference, we also purified translocon mutant forms containing three combinations of Cys-to-Ser pair substitutions in LptD (LptD^CCSS^, LptD^CSCS^ and LptD^SCSC^). With the overproduction of LptM, nickel-affinity purification of LptE^His^ yielded approximately 95% of LptD in its mature oxidized state, LptD^Ox^ (Figure 3D, lane 2). In contrast, a significantly lower amount of LptD^Ox^ was obtained in the absence of LptM (Figure 3D, lane 1). In this sample, approximately one third of LptD was either reduced or only partially oxidized (Figure 3D, quantification). These LptD oxidation intermediate forms included LptD^C1-C2^ (migration similar to that of LptD^CCSS^) and a LptD form containing a non-consecutive disulfide bond that migrates slower than LptD^Ox^, seemingly corresponding to LptD^C2-C4^ (Figure 3D, compare lane 1 to lanes 4 and 5). We also found that the complex containing LptD^CCSS^ migrates slower on BN gels compared to the complex containing wild-type LptD (Figure 3E, lanes 1 and 2). These results suggest that, compared to the LPS translocon containing LptD^Ox^, the presence of LptD^C1-C2^ retards the complex migration on BN gels, providing a plausible explanation for the differential BN-PAGE separation patterns of LptDE purified from wild-type cells (containing mostly LptD^Ox^) or from Δ*lptM* cells (containing also partially oxidized LptD forms) (Figures 1A, lanes 3 and 4, and 3E, lanes 3 and 4). We conclude that LptM facilitates LptD oxidative maturation, whereas its inactivation causes the accumulation of intermediate oxidation forms, most prominently LptD^C1-C2^ (Figures 3B and 3C).

### LptM functions synergistically with DsbA during LptD oxidative maturation

The observation that Δ*lptM* accumulates the LptD^C1-C2^ oxidation intermediate suggests that LptM is not strictly required for DsbA activity. Yet LptM could have a different role in LptM oxidative maturation, thus acting synergistically with DsbA. To test this hypothesis, we generated a double deletion strain lacking *lptM* and the cysteine oxidase *dsbA* by using a P1 phage transduction protocol. On LB agar plates, the Δ*lptM DdsbA* strain formed slightly smaller colonies compared to its parental single deletion strains (Figure 4A, left). Furthermore, the double deletion strain showed an enhanced sensitivity to both vancomycin and SDS (Figure 4A, center and right), indicative of a pronounced OM permeability barrier defect that was not observed with the single deletion strains. This result implies that, as hypothesized, LptM acts synergistically with DsbA in promoting proper oxidative maturation of LptD.

**Figure 4.**
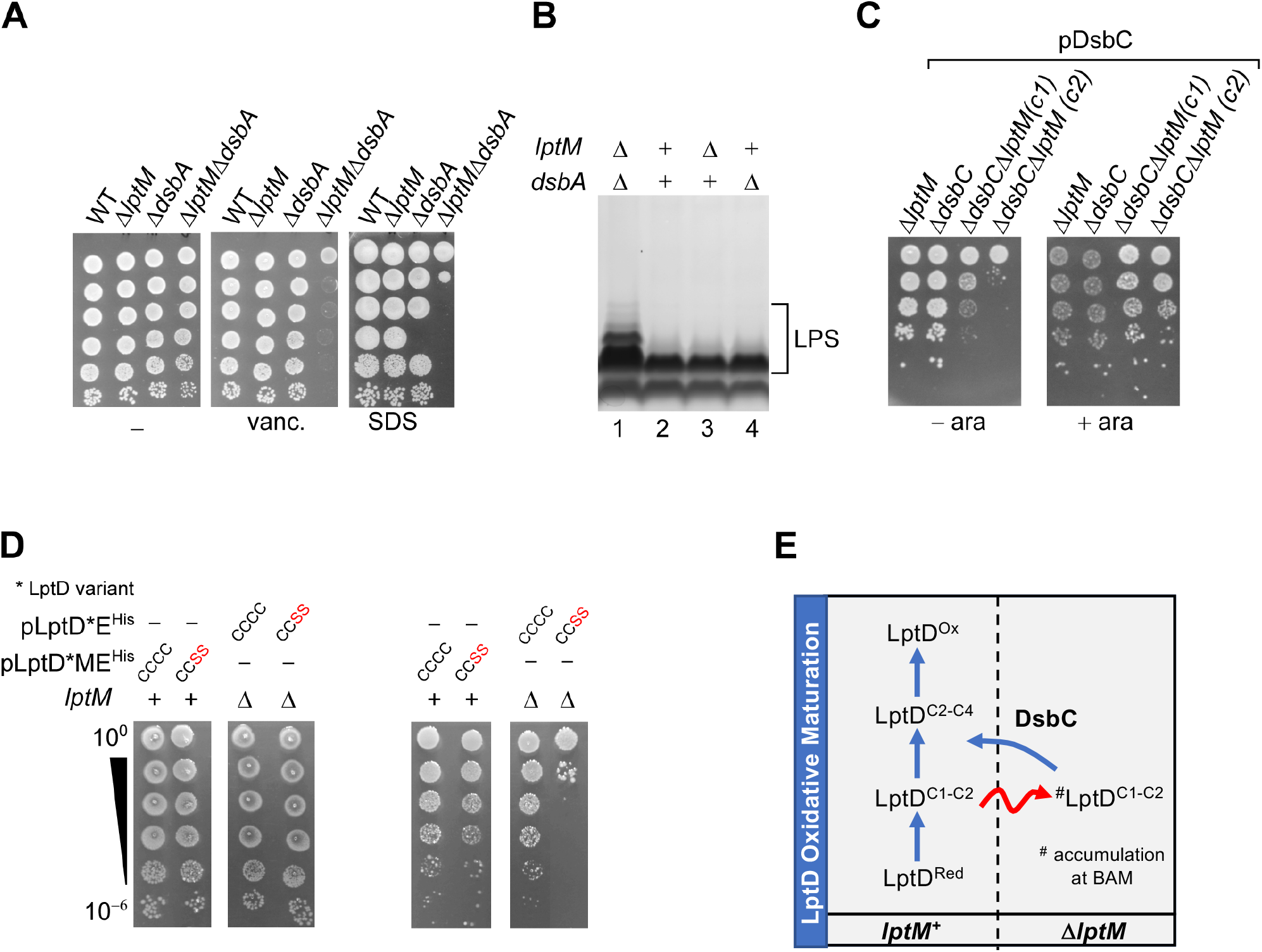
LptM functions synergistically with the oxidative folding machinery. **(A)** Drop dilution growth test of *E. coli* wild-type and the indicated derivative strains lacking *lptM, dsbA* or both under different conditions as indicated. **(B)** LPS was extracted from the indicated strains, resolved by SDS-PAGE and silver stained. **(C)** Drop dilution growth test of the indicated derivative strains lacking *lptM, dsbC* or both transformed with pDsbC on media lacking or supplemented with 0.02% arabinose. **(D)** Drop dilution growth test of *lptM^+^* or Δ*lptM* strains transformed with the indicated plasmids pLptD*E^His^ or pLptD*ME^His^ on media lacking or supplemented with 100 μM IPTG. The “ * “ on the name of the plasmid indicates potential variants of plasmid-encoded LptD, wild-type LptD^CCCC^, or LptD^CCSS^, as specified on the top of each dilution test. **(E)** Schematic representation of the LptD oxidative maturation pathway. Left: In wild-type cells (*lptM^+^*), the four cysteines of LptD are oxidized stepwise: LptD^Red^ is first oxidized to form LptD^C1-C2^. Disulfide bond shuffling in the latter generates LptD^C2-C4^. A final event of oxidation forms a disulfide between C1 and C3, thus generating LptD^Ox^. LptM acts synergistically with the disulfide bond formation machinery in facilitating proper oxidation of LptD. Right: In the absence of LptM, a considerable fraction of LptD accumulates as oxidation intermediate forms, most prominently LptD^C1-C2^. In Δ*lptM*, assembly of LptD together with LptE stalls at the BAM complex (#LptD^C1-C2^). In this strain, DsbC is essential for cell viability, suggesting that disulfide bond isomerization in #LptD helps to rescue LPS translocon assembly.

We also investigated whether the detrimental effect caused by the *lptM dsbA* double deletion was related to impaired LPS transport. LPS extracted from the wild-type and the single deletion strains showed similar SDS-PAGE and silver staining patterns (Figure 4B, lanes 2-4), whereas slower migrating forms of LPS were obtained from the double *lptM dsbA* deletion strain (Figure 4B, lane 1). A mass increment of LPS reveals a modification that can be triggered as a consequence of impaired anterograde transport along the Lpt pathway, causing LPS accumulation at the IM and modification with colanic acid (Meredith et al., 2007; Sperandeo et al., 2008). Most importantly, the observed LPS modification in the *lptM dsbA* double deletion strain provides direct evidence that LptM promotes LPS translocon biogenesis, thereby facilitating efficient LPS transport to the OM.

### In the absence of LptM, DsbC is crucial for cell viability

To gain further insights into how LptM functions together with the oxidative folding machinery, we sought to generate the double deletion strains Δ*lptM dsbC::kan* or Δ*dsbC lptM::kan*. However, we could not combine the deletion of *lptM* with that of *dsbC* by using P1 phage transduction (see Methods for data quantification), suggesting that *lptM* is synthetically lethal with *dsbC*. Indeed, a strain lacking both chromosomal *dsbC* and *lptM*, and encoding DsbC under the control of an arabinose-inducible promoter, *P_BAD_*, could grow efficiently only in the presence of the inducer (Figure 4C), indicating that *dsbC* and *lptM* are nearly essential upon inactivation of the other. This result shows that in the absence of LptM, LptD disulfide bond isomerization by DsbC becomes critical.

To gain deeper understanding of the consequences related to impaired LptD disulfide bond isomerization, we challenged cells with the overproduction of LptE and wild-type LptD (LptD^CCCC^) or LptD^CCSS^, *i.e*. a form of LptD that can be converted to LptD^C1-C2^ but that cannot undergo subsequent disulfide bond shuffling as it contains serine residues in place of C3 and C4. Notably, the overproduction of LptD^CCSS^ impaired growth of Δ*lptM* but not of cells overproducing also LptM (Figure 4D), indicating that the accumulation of LptD^C1-C2^ is detrimental in the absence of LptM. Given the prolonged residence of LptD and LptE at the BAM complex in Δ*lptM* cells (Figure 2B) and that LptD^C1-C2^ is the LptD oxidation intermediate that engages with the BAM complex (Chng et al., 2012; Lee et al., 2016b), it is reasonable to infer that the detrimental effect caused by LptD^C1-C2^ overproduction in Δ*lptM* is related to reduced BAM functionality, which is essential for viability. Together with the observed crucial character of DsbC in Δ*lptM*, our results suggest that in this strain DsbC may help preventing excessive accumulation of LptD^C1-C2^ at the BAM complex (Figure 4E).

### LptM interacts with the OM embedded portion of the LPS translocon

The structures of LptDE from different organisms has been determined (Botos et al., 2016; Botte et al., 2022; Dong et al., 2014; Qiao et al., 2014). To investigate the structural basis of how LptM interacts with LptDE, we used AlphaFold2-multimer to build a predicted model of the complex (Evans, 2022; Jumper et al., 2021; Mirdita et al., 2022). The top-ranking structures reveal a considerable amount of contact between LptM and LptDE (Figure 5A), with the N-terminal cysteine of LptM consistently placed in close proximity of the flexible hinge separating the periplasmic and membrane domains of LptD (Botos et al., 2016; Lundquist and Gumbart, 2020) that is adjacent to the LptD β-barrel lateral gate (Figure S5A), which is consistent with our crosslinking data (Figure S4A and B).

**Figure 5.**
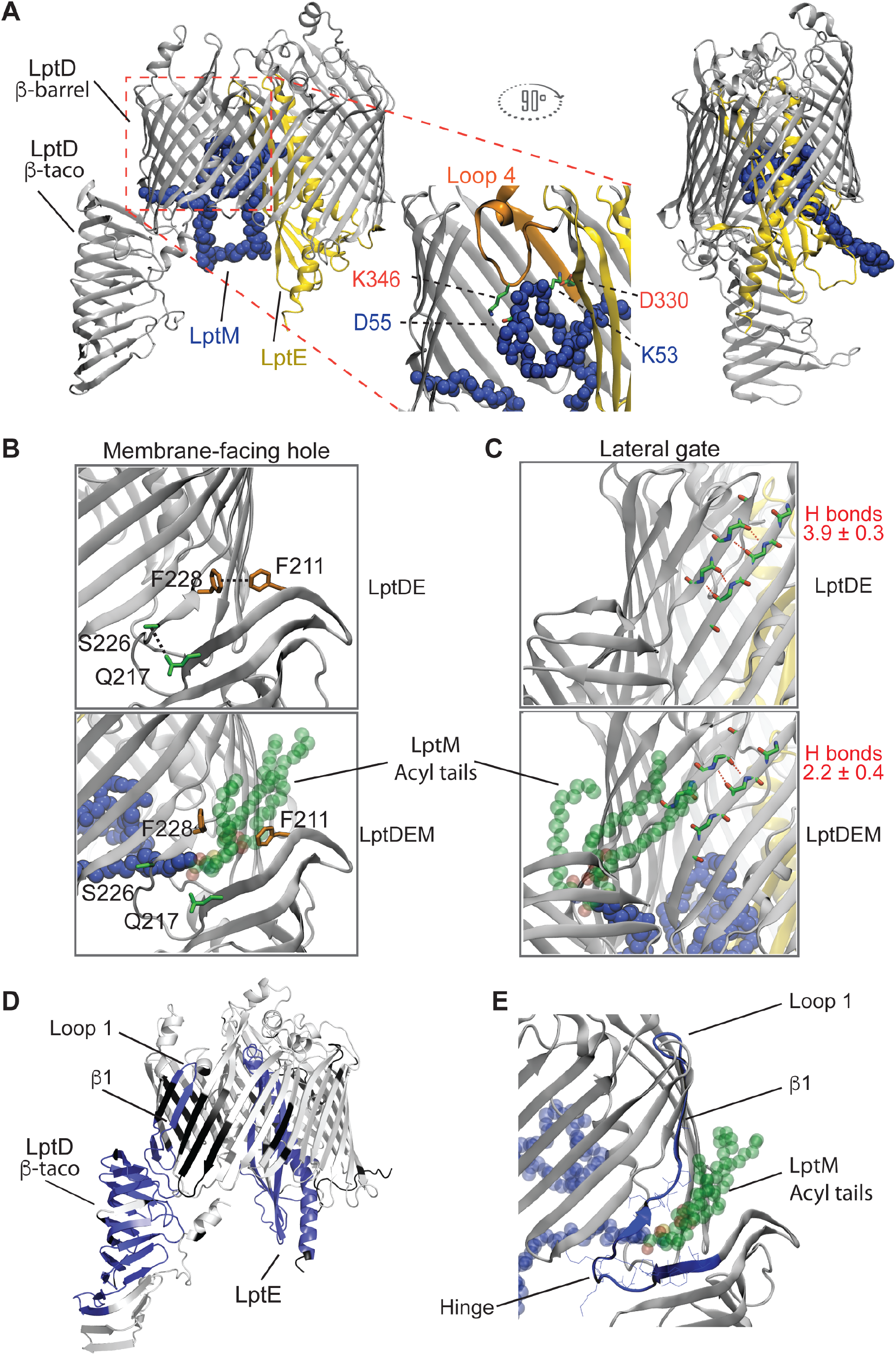
LptM binds the membrane-embedded portion of the LPS translocon, influencing LPS-interaction sites. **(A)** View of top ranking AlphaFold2 model of LptDE-LptM. LptD is shown as gray cartoon, LptE as yellow cartoon, and the LptM backbone is shown as blue spheres. A zoom-in shows the interactions of LptM with the LptD Loop 4 (orange cartoon). Salt-bridges predicted in the AlphaFold2 model are shown. **(B)** Zoom in on the LptD hinge between the β-taco and β-barrel in either the LptDE (top) or LptDE-LptM (bottom) systems. Image from a snapshot following 500 ns of MD simulation. Several interactions are made between the taco and barrel in LptDE but broken when LptM is present (blue and green spheres), including between the residues shown. **(C)** View of the LptD lateral gate with quantification of H-bond number from 3 x 500 ns simulations for the LptDE and LptDE-LptM systems. Representative H-bonds are shown as dashed red lines, as computed using VMD. **(D)** Differential deuterium uptake between the LptDE translocon in the presence or absence of LptM after statistical analysis with Deuteros, showing in blue regions that are significantly protected upon LptM binding. **(E)** View of the AlphaFold2 model plus modelled LptM tri-acylation, showing the position of the LptM N-terminus and tri-acylation (blue and green transparent spheres) in relation to the LptD 212-240 peptide, which is protected from deuteration by LptM in the HDX-MS experiments. Residues from the 212-240 peptide which are in contact with LptM (<0.3 nm) are shown as lines.

We next investigated how LptM might influence the conformational dynamics of the OM LPS translocon using atomistic molecular dynamics (MD) simulations of our AlphaFold2-multimer model. The presence of LptM N-terminus close to the LptD hinge was compatible with C1-C3 and C2-C4 being in disulfide bond distances over a simulation time of 500 ns, whereas C1 and C2 remained more distant from each other (Figure S5A). Furthermore, the positioning of the N-terminal cysteine means that, when tri-acylated, the acyl tails are positioned at the interface of the β-taco and β-barrel domains (Figure 5B). This region presents several hydrophobic residues and was proposed to form a membrane-facing hole through which the acyl chains of LPS would get inserted into the OM (Gu et al., 2015). From our simulations, the presence of LptM at this position breaks several connections of the β-taco with the β-barrel domain (Figure 5B), which has an impact on the angle of orientation of the LptD β-taco domain (Figure S5B) and causes destabilization of the N-terminal loop of LptD that precedes the β-taco domain (Figure S5C). LptM also affects the LptD β-barrel lateral gate, reducing the number of H-bonds formed between β-strands 1 and 26 at their periplasm-facing side (Figure 5C). This is in line with a previously proposed model of lateral gate opening (Botte et al., 2022) and suggests that LptM may reduce the energetic barrier of lateral gate opening (Dong et al., 2014; Lundquist and Gumbart, 2020).

Aside from these interactions, our model also predicts that LptM inserts into the lumen of the LptD β-barrel, establishing several interaction points with both LptD and LptE. These include a salt bridge formed between LptM K23 and LptD E275 (Figure S5D). Our model further predicts that the C-terminal region of LptM forms a hook structure that gets positioned between the inward folded LptD loop 4 and LptE (Figure 5A, frame). More specifically, in our model and simulations, we observed high degrees of contact between LptM and the LptD segment D330-D352, which includes part of β 7 and loop 4. In fact, salt bridges were predicted between LptM K53 and LptD D330 as well as LptM D55 and LptD K346 (Figure 5A, frame). This result is particularly intriguing as the phenotype caused by the deletion LptD330-352 (encoded by the mutant allele *lptD4213*) resembles that of Δ*lptM*, that is impaired OM LPS translocon assembly, stalling of the LPS translocon at the BAM complex and accumulation of the oxidation intermediate LptD^C1-C2^ (Chng et al., 2012; Lee et al., 2019; Lee et al., 2016b).

Our structural model suggests that LptM mostly interacts with the OM-embedded portion of the LPS translocon. To verify this prediction, we purified LptM^His^ in cells that express, in addition to endogenous wild-type LptD, also a form of LptD consisting of the β-barrel domain that is deleted of the N-terminal β-taco domain and that can still interact with LptE (Botos et al., 2016; Dong et al., 2014; Qiao et al., 2014). In line with our AlphaFold2-multimer model, we found that the lack of the β-taco domain does not influence the interaction of LptM with truncated LptD and LptE (Figure S6).

We also employed hydrogen-deuterium exchange mass spectrometry (HDX-MS) to monitor the dynamics and solvent accessibility of LptD and LptE in the LPS translocon produced in the presence or absence of LptM and purified in detergent micelles. We obtained 93% and 75% sequence coverage for LptD and LptE, respectively, showing that the deuteration was very slow for the membrane-spanning portion of the LptD β-barrel domain and much faster in its periplasmic region, and extracellular loops (Figure S7), as expected for an integral membrane protein (Donnarumma et al., 2018; Javed et al., 2022). Less expected was the relatively high deuteration rates obtained for LptE, as this protein buries in the internal lumen of the LptD β-barrel. Comparing the deuteration of the LPS translocon with and without LptM, we observed that LptM stabilizes distinct regions of the OM LPS translocon in a statistically significant manner (Figures 5D, S8 and S9). These stabilized regions include LptE (Figures 5D, S8, S9) and a segment of LptD (212-240) corresponding to the hinge followed by β 1 and loop 1 of the β-barrel domain (Figures 5E, S8, S9, and S10). The protection of LptE is in line with the extensive contacts between LptM and LptE in the lumen of the LptD β-barrel domain predicted by our AlphaFold2-multimer model. Similarly, protection of the β-barrel N-terminal peptide (LptD 212-240) fits well with the predicted positioning of the LptM N-terminus in proximity of the LptD β-taco/β-barrel hinge and the β-barrel lateral gate. This segment would also be proximal to the acyl tails of the LptM N-terminal cysteine, which might contribute to its stabilization (Figures 5E and S9). Notably, a recent HDX-MS analysis of *Klebsiella pneumoniae (Kp*) LptDE showed that, upon LPS binding, deuteration of the N-terminal region of the LptD β-barrel in proximity to the lateral gate is reduced (Fiorentino et al., 2021). Thus, the presence of LptM produces a stabilization effect in proximity of the LptD lateral gate that is similar to that caused by LPS binding. Taken together, our structure modeling and HDX-MS results suggest an LPS-mimicking effect by LptM in the membrane embedded portion of the LPS translocon.

Deuteration of the β-taco domain was also decreased in the presence of LptM. A bimodal isotopic distribution (EX1 regime) for peptides of the β-taco domain was previously reported for *Kp* LptDE (Fiorentino et al., 2021). Based on this result, it was suggested that the β-strands of the β-taco domain undergo concerted closing and opening motions that occur with a kinetic slower than the rate of hydrogen/deuterium exchange. For peptides of the central and C-terminal region of the β-taco domain, LPS binding favored a mixed kinetics of deuterium uptake, whereas the antimicrobial peptide thanatin, which also binds the LptD β-taco domain (Vetterli et al., 2018), shifted the equilibrium toward a slower uptake kinetics (Fiorentino et al., 2021). We observed a similar bimodal isotopic distribution for peptides of the β-taco (LptD 70-218) of the LptDE dimer in the absence of ligands, as previously reported (Fiorentino et al., 2021) (Figure S10). Strikingly, the presence of LptM abrogated this bimodal regime and shifted the mixed kinetics equilibrium toward the slower uptake behavior (Figures 5D, S8 and S10), suggesting an effect similar to that shown for *Kp* LptDE in the presence of thanatin. Taken together, these results provide strong evidence that the extensive contacts predicted between LptM and distinct regions of the LPS translocon may have an important regulatory role.

## DISCUSSION

LPS transport was known to be mediated by seven proteins, LptA-G (Okuda et al., 2016). Here we report the discovery and characterization of an eighth component of the LPS transport machinery, LptM. We have shown that the lipoprotein LptM stably interacts with the OM-embedded portion of the LPS translocon formed by LptD and LptE, promoting its assembly by the BAM complex. Our functional analysis of LptM provides novel insights into the mechanisms that activate the LPS translocon upon assembly by the BAM complex and clarifies the role of disulfide bond isomerization by DsbC during oxidative maturation of LptD. Furthermore, the biochemical and structural analysis of the LptDEM heterotrimer highlights how several translocon domains reported to be crucial in coordinating LPS transport are influenced by LptM.

In this study, we provide three major lines of evidence demonstrating the critical role of LptM in the assembly of the OM LPS translocon. First, LptM interacts with the LptD β-barrel domain and LptE. This finding is fully supported by our biochemical assays, native-MS analysis, AlphaFold2 modeling, atomistic MD simulations and HDX-MS kinetics. Second, LptM improves the efficiency of LPS translocon assembly at the BAM complex. Third, LptM is required for proper oxidation of LptD and efficient anterograde transport of LPS. Our AlphaFold2-multimer model and atomistic simulations provide important clues as to how LptM promotes LptD maturation. LptM stably associates via both its N- and C-terminal regions with the membrane-embedded portion of the LPS translocon. This association is mediated by several electrostatic interactions, including two salt bridges that connect LptM to a segment of LptD (residues D330-D352), comprising part of β 7 and of the inward-folded extracellular loop 4 of the LptD β-barrel domain (Botos et al., 2016; Dong et al., 2014; Qiao et al., 2014). This segment is critical for LptD assembly as a mature LPS translocon, whereas its deletion causes a partial loss of function (Sampson et al., 1989; Wu et al., 2005). Deletion of LptD D330-D352, encoded by the well-characterized allele *lptD4213*, increases the time of residence of LptD at the BAM complex causing the accumulation of the non-functional LptD^C1-C2^ assembly intermediate (Chng et al., 2012; Lee et al., 2019). The LptD assembly defect caused by the deletion of *lptM* (prolonged residence at the BAM complex and impaired oxidative maturation) is particularly reminiscent of that previously reported for the *lptD4213* allele. This observation suggests that the interaction of LptM with LptD loop 4 may facilitate the correct positioning of this surface exposed region that is critical for the progression of LptD maturation to form a fully oxidized and functional LPS translocon.

Formation of native disulfide bonds in LptD is carefully controlled and occurs only after the β-barrel domain of LptD engages with the BAM complex to assemble together with LptE into the OM (Chng et al., 2012; Lee et al., 2016b). Despite the fact that native disulfide bonds are crucial for LPS translocon activation, DsbA is not essential for viability indicating that spontaneous oxidation sustains formation of a sufficient amount of properly oxidized LptD (Meehan et al., 2017). Our results indicate that, with the inactivation of the oxidative folding machinery, LptM is particularly critical for efficient LPS translocon activity. Thus, LptM acts independently of but synergistically with the oxidative folding machinery in promoting LPS translocon activation. As indicated by our structural modeling and HDX-MS analysis, LptM binds and stabilizes regions of the OM-embedded portion of the translocon. By doing so, it is plausible that LptM stabilizes a conformation of the translocon that can most effectively be oxidized to form native disulfide bonds, even in the absence of a functional oxidative folding machinery (Figures 4A and 4B).

Our results show that LptM is not required to generate the LptD^C1-C2^ oxidation intermediate, as this accumulates in the absence of LptM (Figures 3B and 3D). Instead, LptM plays an important role after this first LptD oxidation event mediated by DsbA. Different views exist on the mechanisms that mediate disulfide bond isomerization in LptD^C1-C2^. Experimental evidence that DsbC can form an intermolecular disulfide bond with LptD suggested that this enzyme could be involved in the rearrangement of its cysteine oxidation patterns (Denoncin et al., 2010). Yet, the observation that DsbC is not essential for cell viability casted doubts on its role during functional activation of the LPS translocon (Ruiz et al., 2010). Our results clearly indicate that DsbC is crucial for viability in cells that lack LptM. In these cells, LptD and LptE reside at the BAM complex for a longer period of time compared to cells that express LptM, which is indicative of a stalled assembly process. Stalling of a LptD assembly intermediate at the BAM complex may potentially have a catastrophic effect on the OM as it will interfere not only with LPS translocon biogenesis but also with the ability of the BAM complex to assemble other proteins into the OM. The latter scenario is supported by the fact that overproduction of LptD^CCSS^, in which the disulfide bond C1-C2 cannot undergo isomerization, is particularly detrimental in D*lptM*. Our results imply that DsbC plays a role in rescuing the stalled LptD^C1-C2^ intermediate (Figure 4E), thus preventing a catastrophic effect on the OM. Taken together, we conclude i) that LptM acts downstream of the first oxidation event mediated by DsbA and ii) that, in the absence of LptM, LptD oxidation intermediates that fail to assemble into an active translocon require rescuing by DsbC (Figure 4E). These considerations support a mechanistic model where LptM stabilizes a conformation of the LPS translocon in which LptD can efficiently complete oxidative maturation.

Finally, we have obtained evidence that LptM, besides being crucial for the correct assembly of the LPS translocon into the OM, may also regulate its function. LptM binds the periplasm-facing luminal region of the plug-and-barrel translocon structure. The N-terminus of LptM interacts with a region of the translocon in proximity of the LptD lateral gate. Intriguingly, our structural model shows that the acyl tails at the N-terminus of LptM would insert into the membrane-facing hole that forms at the interface between the β-taco and the β-barrel domains of LptD reducing the hydrogen-bonding of the lateral gate. This result was validated by our HDX-MS analysis, showing that LptM reduced deuteration of the LptD lateral gate. The stabilization effect of LptM on the lateral gate of LptD is similar to that of LPS-binding observed for *Kp* LptD (Fiorentino et al., 2021). The interaction of the LptM C-terminal region with LptD loop 4 also points to a critical regulation of the LPS translocon, as this loop closes the lumen of the barrel to the extracellular space and has been proposed to guide the polysaccharide chain of LPS into the external leaflet of the OM (Botos et al., 2016; Dong et al., 2014; Li et al., 2015; Qiao et al., 2014). Similarly, LptM interacts extensively with LptE, which also binds LPS molecules (Malojčić et al., 2014). Thus, LptM binds region of the translocon that are crucial to coordinate transport of LPS into the OM. We propose that, by mimicking LPS binding in the membrane-embedded region of the translocon, LptM stabilizes its active conformation. Importantly, whereas LptM appears to mimic LPS binding in the OM-embedded portion of the LPS translocon, it also rigidifies the periplasmic β-taco domain of LptD, suggesting tight control of the initial event of LPS docking onto the translocon (Fiorentino et al., 2021). We can postulate that, during LPS transport, the initial event of LPS binding to the β-taco domain increases translocon dynamics and progressively promotes LPS insertion into the OM via the LptD membrane-facing hole and lateral gate. A possible scenario is that LPS breaches the OM by a mechanisms of substrate exchange, where LPS substitutes LptM.

The broad conservation of the N-terminal LptM PF13627 protein motif in proteobacteria and, most notably, the conservation of full-length LptM in the *Enterobacteriaceae* family, suggest that the newly identified role of LptM in LPS translocon biogenesis can be generalized to other bacteria. The two essential surface-exposed machineries, the OM LPS translocon and the BAM complex are emerging as attractive targets for the development of novel antimicrobial compounds (Robinson, 2019; Sousa, 2019). In this context, the newly described mechanism of LPS translocon binding by the small lipoprotein LptM can help identifying new target sites in LptDE. More generally, by discovering LptM as a novel LPS translocon component that is assembled together with LptD and LptE at the BAM complex, our study provides a key piece of information that will be of paramount importance in developing OM-targeting drugs.

## METHODS

### Bacterial strains and growth conditions

The *E. coli* strains used in this study are listed in Table 1. All strains are derived from BW25113 (Grenier et al., 2014). Gene deletions were obtained by P1 phage transduction using a P1 phage lysate of the corresponding Keio collection strain (Baba et al., 2006). Double gene deletions were obtained by performing a second P1 phage transduction after excising the kanamycin-resistance cassette from the first mutated locus using the heat-curable plasmid pCP20 (Datsenko and Wanner, 2000). Transduction of wild-type and Δ*lptM* cells with a Δ*dsbC* P1 phage lysate gave rise respectively to 137 and 0 colonies, respectively, whereas wild-type and Δ*dsbC* cells supplemented with the Δ*lptM* P1 phage lysate gave rise to 570 and 0 colonies, respectively. Thus, to build a *dsbC lptM* double deletion strain, the Δ*lptM* strain was first transformed with pDsbC, harbouring *dsbC* under the control of the arabinose-inducible P_*BAD*_ promoter. This *dsbC* diploid strain was subjected to P1 phage transduction to delete chromosomal *dsbC*. The produced DsbC depletion strain was grown in the presence of 0.02% arabinose. *E. coli* strains were cultured on lysogenic broth (LB) liquid media or LB agar plates supplemented with the following antibiotics: 100 μg/ml ampicillin, 50 μg/ml kanamycin, 30 μg/ml chloramphenicol. Serial dilution assays were conducted on LB agar plates supplemented with 5 μg/ml or 60 μg/ml vancomycin, 0.2% (w/v) SDS, or 100 μM Isopropyl β-D-1-thiogalactopyranoside (IPTG), as indicated.

**Table 1.**
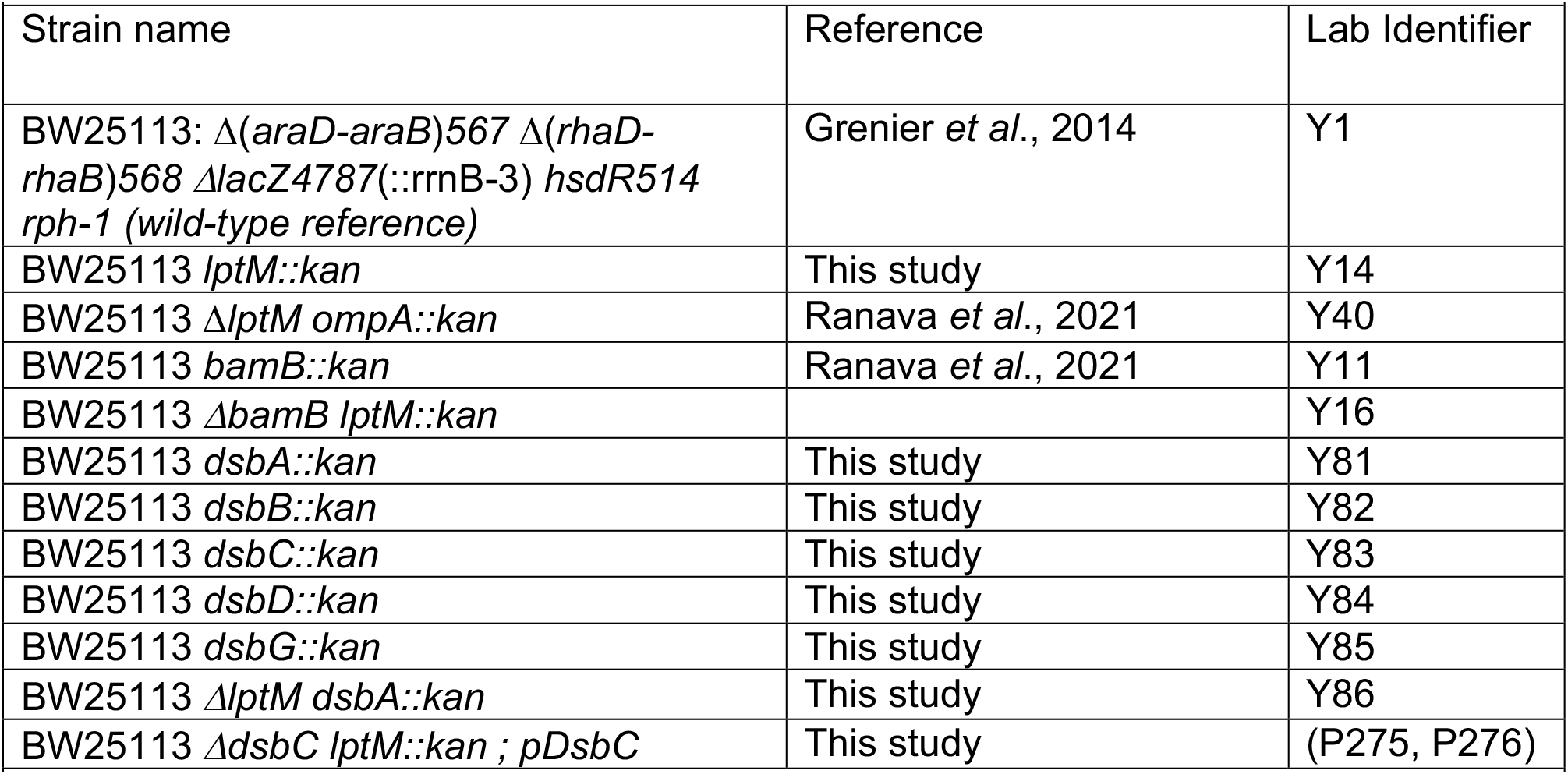
Bacterial strains.

### Plasmid construction

Plasmids and oligonucleotides are listed in Tables 2 and 3, respectively. Plasmid pDsbC was obtained by cloning *dsbC* under an arabinose inducible promoter in the pBAD33 vector (Guzman et al., 1995). All other plasmids were derived by cloning genes of interest downstream of an IPTG inducible promoter in the pTrc99a vector. pLptM^His^, pLptDE^His^, pLptDME^His^, pLptDEM^His^ and pDsbC were constructed by overlap extension PCR cloning. Site-directed mutagenesis was used to replace specific codons in pLptM^His^ with an amber codon for incorporation of the unnatural amino acid analog *p*-benzoyl-L-phenylalanine (pBpa) by amber suppression (Chin et al., 2002). Site-directed mutagenesis was also used to replace cysteine-with serine-encoding codons in pLptDE^His^ or pLptDME^His^. pLptD^β-barrel^EM^His^ encoding a version of LptD that lacks amino acids 25-205 (corresponding to the β-taco domain) was constructed by inverse PCR on pLptDEM^His^.

**Table 2.**
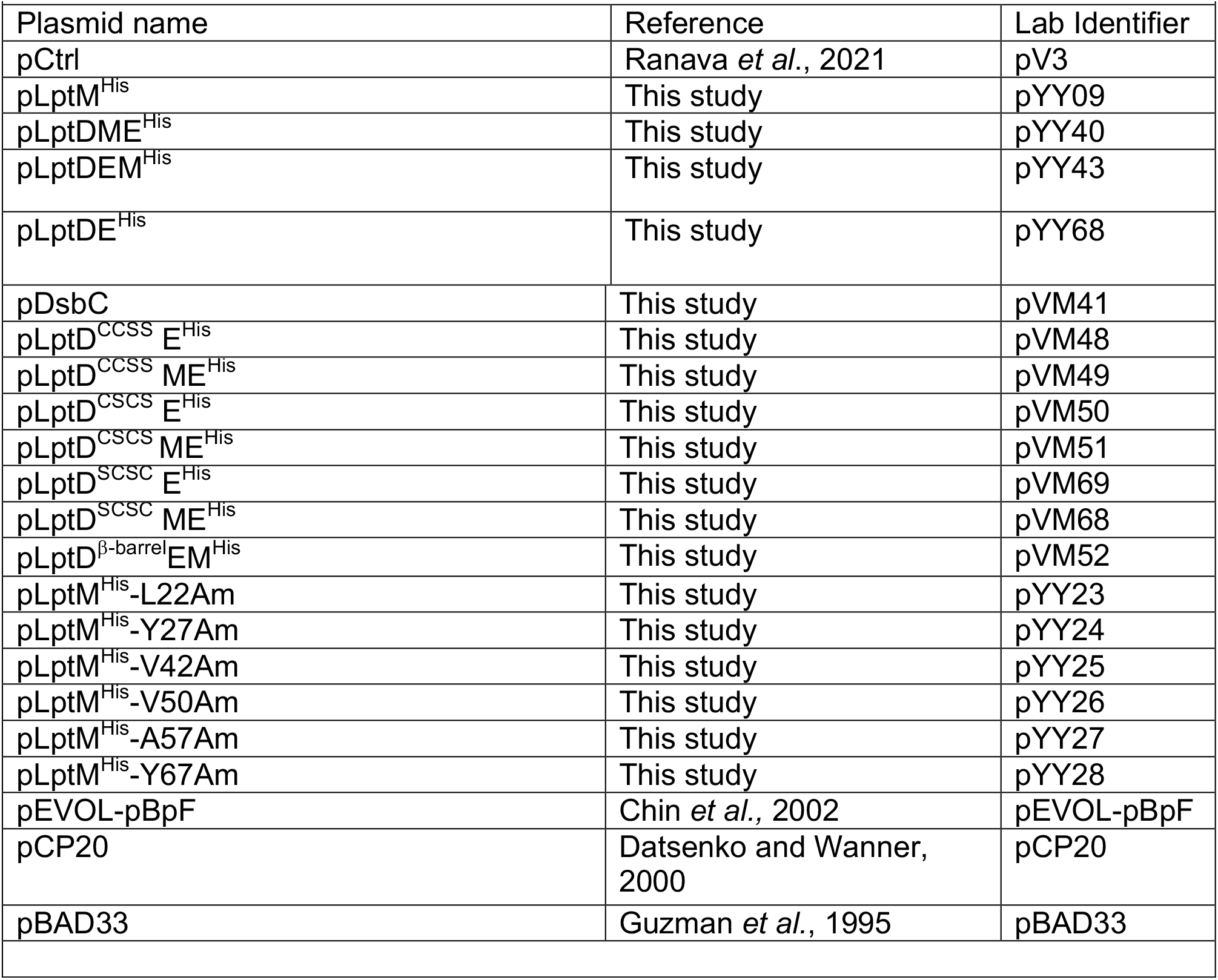
Plasmids.

**Table 3.**
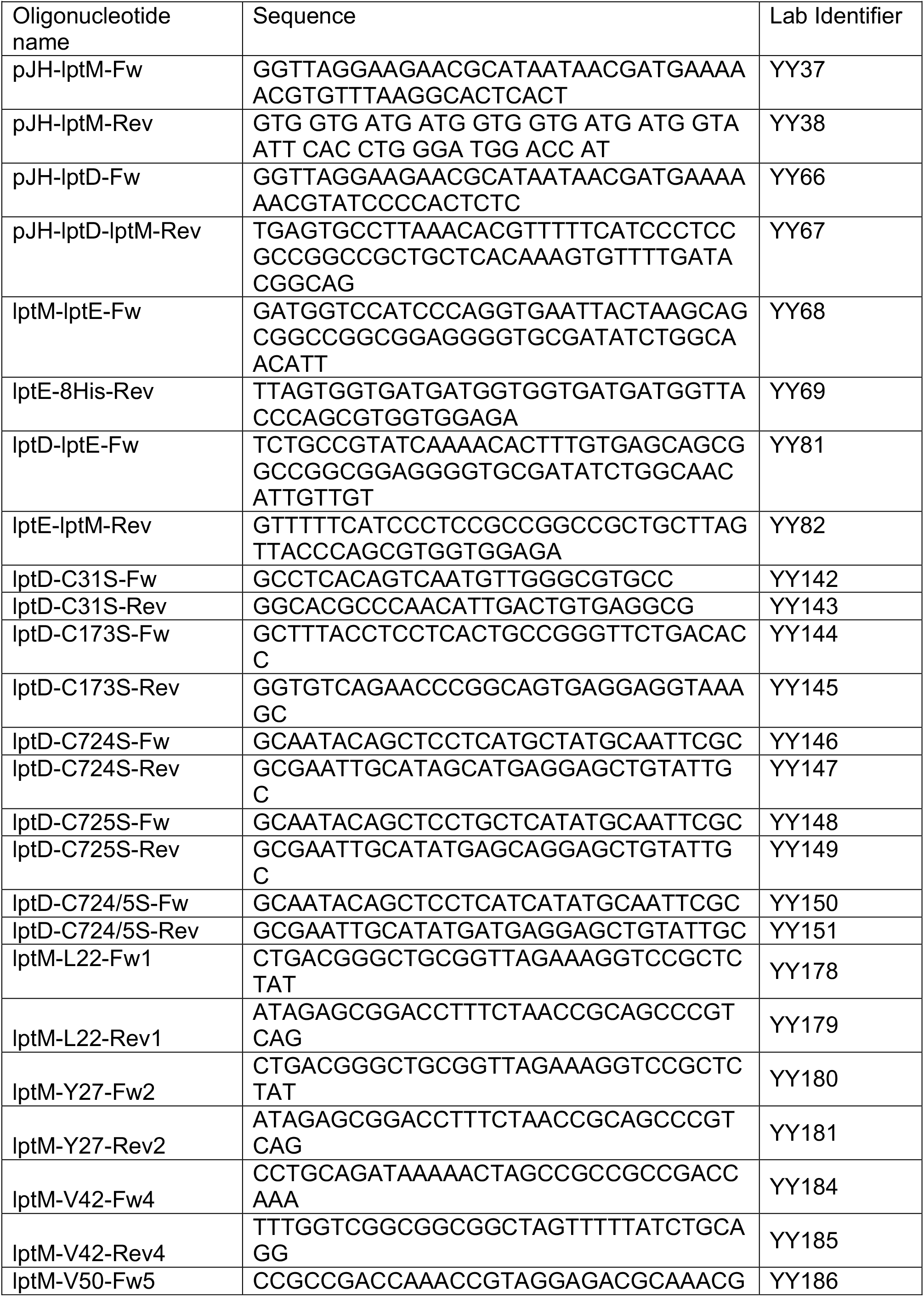
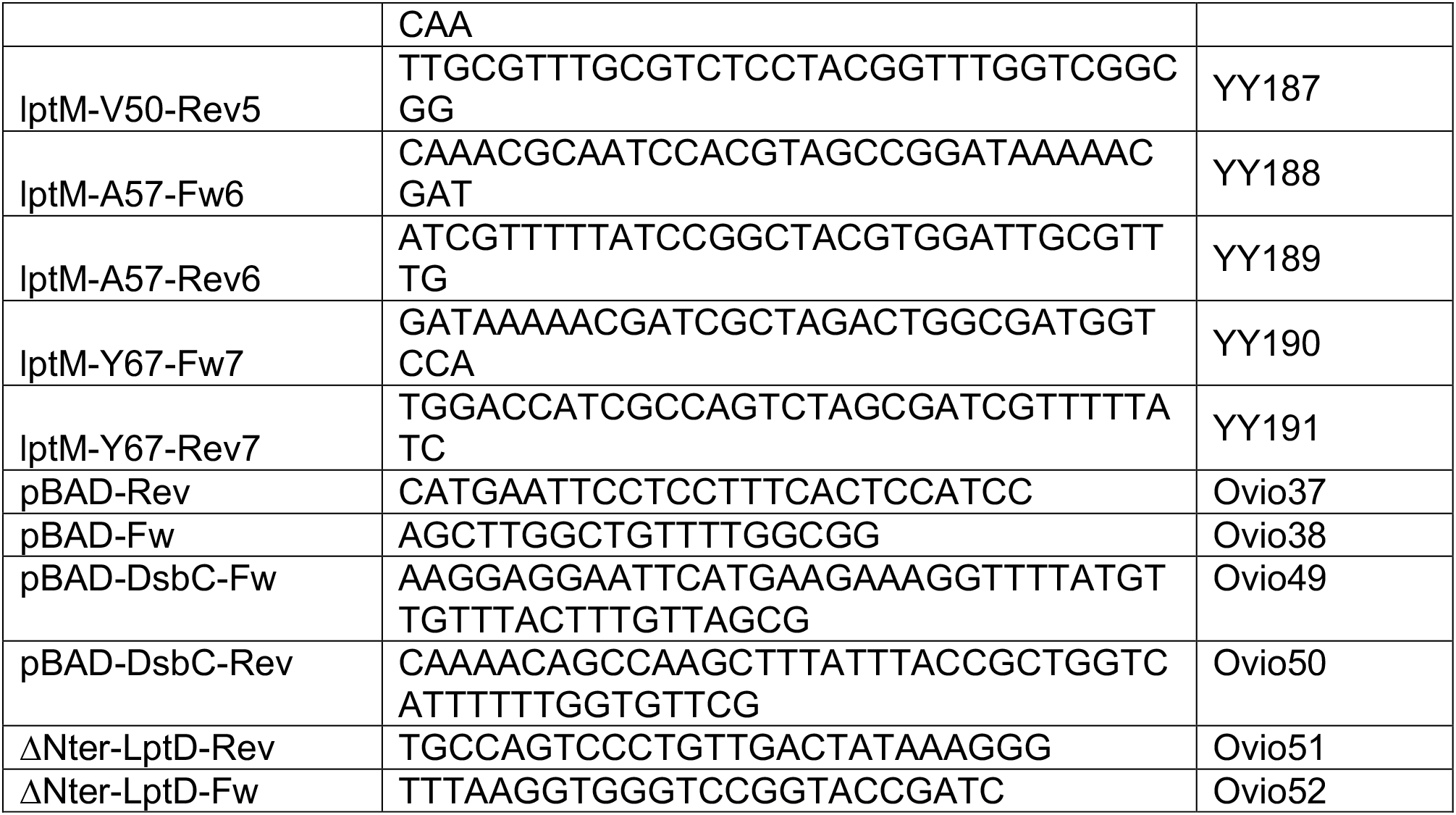
Oligonucleotides.

### Taxonomic analysis of LptM

The procedure for the taxonomic analysis of LptM is detailed in the Methods section of the Supplementary Information file. The logoplot of the multiple alignment of LptM amino acid sequences was obtained using ggseqlogo (Wagih, 2017).

### Cell fractionation and isolation of native protein complexes

Whole cell lysate was prepared from cells cultured to mid-exponential phase at 37°C. Where indicated, cells were supplemented with IPTG for 1.5h prior to collection. Cells were pelleted by centrifugation, lysed in Laemmli sample buffer (Bio-Rad) and subjected to boiling. Protein affinity purification under native conditions was conducted as previously described (Ranava et al., 2021). Briefly, when the cell cultures reached mid-exponential phase (OD600=0.5) protein expression was induced by supplementing 200 μM IPTG for 1.5h prior to cell collection by centrifugation. Cells were resuspended in 20 mM Tris-HCl pH 8 containing a EDTA-free protease inhibitor cocktail (Roche). For the purification of protein samples that had to be analyzed both by reducing and non-reducing SDS-PAGE, the cell resuspension buffer was further supplemented with 50mM iodoacetamide (Sigma) to alkylate free thiol groups of Cys residues. Resuspended cells were mechanically disrupted using a cell disruptor (Constant Systems LTD) set to 0.82 kPa. The obtained cell lysate was clarified by centrifugation at 6,000 x *g*, 4°C for 15 min. The crude envelope fraction was collected by subjecting the supernatant to ultracentrifugation at 50,000 x *g* at 4 °C for 30 min. To perform affinity purification of protein complexes, the crude envelope fraction was solubilized with 50mMTris-HCl pH 8, 150mM NaCl, 20 mM imidazole supplemented with EDTA-free protease inhibitor (Roche) and 1% (w/v) n-dodecyl-β-D-maltopyranoside (DDM, Merck) or 1% (w/v) digitonin (Merck). After a clarifying spin to remove insoluble material, solubilized proteins were incubated with Protino Ni-NTA resin (Machery-Nagel) for 2h at 4°C. After extensive washes of the column with 50 mM Tris-HCl pH 8, 150mM NaCl, 50 mM imidazole and 0.03% (w/v) DDM, bound proteins were eluted with a similar buffer contain 800 mM imidazole and 10% (w/v) glycerol. Aliquots of the elution fractions were snap-frozen in liquid nitrogen for storage at −80 °C or directly analyzed by SDS or blue native gel electrophoresis.

### MALDI-TOF/TOF mass-spectrometry

Excised bands from BN-PAGE were washed by 25 mM NH_4_HCO_3_, pH 7.8-acetonitrile 50:50 (v/v), and digested by trypsin. MS and MS/MS experiments were carried out using a-cyano-4-hydroxycinnamic acid matrix at a concentration of 6 mg/ml in 50 % (v/v) acetonitrile-0.1% trifluoroacetic acid. Analyses of trypsin digested samples were performed on a MALDI TOF/TOF, in reflector positive mode. Parameters were set to source and grid voltages to 20 and 14 kV, respectively, power laser from 2000 to 3500, extraction delay time, 200 ns; shoot number, 1000. Acquisition range was between 800 and 3500 m/z. Spectra were analyzed using Data Explorer software (Applied Biosystems). MS/MS spectra were acquired using a MS/MS positive acquisition method, with 1 kV positive operating mode, and a CID off mode. The MS/MS spectra were examined and sequenced based on assignment of the N-terminal b-ion and C-terminal y-ion series.

### Native Mass Spectrometry

Prior to native MS analysis, 100 μl LptDE^His^ (7 μM) and LptDEM^His^ (10 μM) samples were desalted in 200 mM ammonium acetate, pH 7.4 supplemented with 0.03% DDM using ultrafiltration with MWCO 30 kDa Vivacon 500 (Sartorius) and concentrated to ~10-20 μM. Samples were analyzed on a SYNAPT G2-Si mass spectrometer (Waters, Manchester, UK) running in positive ion mode and coupled to an automated chip-based nano-electrospray source (Triversa Nanomate, Advion Biosciences, Ithaca, NY, USA). The voltages applied to the chip, the sample cone and the ion energy resolving quadrupole were set to 1.8 kV, 200 V and −1.0 V, respectively. Proteins were activation in the collision cell with 200 V trap collision energy and an argon flow of 8 ml/min. The instrument was calibrated with a 2 mg/ml cesium iodide solution in 50% isopropanol. Raw data were acquired with MassLynx 4.1 (Waters, Manchester, UK) and analyzed manually.

### Automated Hydrogen-Deuterium eXchange coupled to Mass Spectrometry (HDX-MS)

HDX-MS experiments were performed on a Synapt-G2 HDMS (Waters Scientific, Manchester, UK) coupled to a Twin HTS PAL dispensing and labelling robot (LEAP Technologies, Carborro, NC, USA) via a NanoAcquity system with HDX technology (Waters, Manchester, UK). Briefly, 5.2 μl of protein at 20 μM were diluted in 98.8 μl of protonated (for peptide mapping) or deuterated buffer (20 mM MES pH 6.5, 200 mM NaCl) and incubated at 20 °C for 0, 0.5, 5, 10 and 30 min. 99 μl were then transferred to vials containing 11 μl of pre-cooled quenching solution (500 mM glycine at pH 2.3). After 30 s quench, 105 μl were injected to a 100 μl loop. Proteins were digested on-line with a 2.1 mm x 30 mm EnzymateTM BEH Pepsin column (Waters Scientific, Manchester, UK). Peptides were desalted for 2 min on a C18 pre-column (Acquity UPLC BEH 1.7 μm, VANGUARD) and separated on a C18 column (Acquity UPLC BEH 1.7 μm, 1.0 mm x 100 mm) by a linear gradient (2% to 40% acetonitrile in 13 min). Experiments were run in duplicates and protonated buffer was injected between each duplicate to wash the column and avoid cross over contamination. Peptide identification was performed with ProteinLynx Global SERVER (PLGS, Waters, Manchester, UK) based on the MSE data acquired on the non-deuterated samples. Peptides were filtered in DynamX 3.0 with the following parameters: peptides identified in at least 3 out of 5 acquisitions, 0.3 fragments per amino-acid, intensity threshold 1000. The Relative Deuterium Uptakes were not corrected for back exchange. Deuteros 2.0 software (Lau et al., 2021) was used for data visualization and statistical analysis. The online web application HDX-Viewer (Bouyssié et al., 2019)and PyMOL (The PyMOL Molecular Graphics System, Version 2.3.0 Schrödinger, LLC) were used to represent the HDX-MS data on either the LptDE and LptDEM models.

The native MS and HDX-MS data has been deposited to the ProteomeXchange Consortium via the PRIDE partner repository (Perez-Riverol et al., 2022) with the dataset identifier PXD038832.

### LPS extraction and silver staining

*E. coli* strains were cultured till OD600 = 0.5. 1.5 ml aliquots of cultures were withdrawn to harvest cells by centrifugation. LPS was extracted following an hot aqueous-phenol extraction (Davis and Goldberg, 2012). Briefly, cell pellets were resuspended in 200 μl blue buffer (50 mM Tris-HCl pH 6.8, 2% (v/v) β-mercaptoethanol, 2% (w/v) SDS, 10% glycerol and a pinch of bromophenol blue). After boiling for 15 min and cooling at room temperature for a similar time, samples were treated with 0.5 mg/ml proteinase K overnight at 59 °C. A first extraction was conducted by applying to the sample 200 μl of ice-cold water-saturated phenol and heating at 65 °C for 15 min with agitation. After cooling to room temperature, 1 ml diethyl ether was applied followed by agitation by vortexing for 15 sec and centrifugation at 20,600 x *g*. The heavier blue LPS-containing phase was withdrawn and approximately 1/10 of the obtained LPS fraction was resolved by SDS-PAGE before direct visualization by Silver staining (SilverQuest Silver Stain, Invitrogen)

### Gel electrophoresis Coomassie staining and Western blotting

Proteins samples were prepared in Laemmli sample buffer (Bio-Rad) lacking β-mercaptoethanol (non-reducing conditions) or supplemented with β-mercaptoethanol (reducing conditions). Proteins were separated by homemade SDS gels (10% acrylamide in Bis-Tris pH 6.4 buffer, subjected to electrophoresis using either MES or MOPS buffer) or home-made blue native 6-13% acrylamide gradient gel (Ranava et al., 2021). Where indicated, gels were stained with Coomassie brilliant blue R250. To perform Western blots, protein gels were blotted onto PVDF membranes (Merck). Upon blocking with skim milk, membranes were incubated with epitope-specific rabbit polyclonal antisera or with an anti poly-histidine horseradish peroxidase-conjugated monoclonal antibody (TaKaRa). Immunodetection was revealed by using a Clarity Western ECL blotting substrate (BioRad) and detected using a LAP-4000 (Fujifilm) apparatus. The signal intensity of protein bands was quantified using the Multi Gauge software (Fujifilm). The rabbit polyclonal antiserum against LptD was a kind gift of Dr. J.F. Collet (UC Louvain, Belgium).

### Protein model building

Models of LptDE and LptDEM were built using AlphaFold2, running AlphaFold Multimer v1.0 via ColabFold (Evans, 2022; Jumper et al., 2021; Mirdita et al., 2022). Models were built using the *E. coli* sequences: residues 26-784 of P31554 for LptD, residues 19-193 of P0ADC1 for LptE, and residues 20-67 of P0ADN6 for LptM (previously YifL). Five rounds were run for each prediction, with the highest ranked model used for follow-up analysis. All AlphaFold2 LptDE and LptDEM models are available for download at https://osf.io/xpfjc/, along with PAE, coverage and plDDT plots. Note that the C-terminus of LptE was truncated after Thr 174 for all images used in the manuscript.

### Site-directed photocrosslinking and purification of crosslinked LptM

The photocrosslinkable amino acid pBpa was introduced at different positions of the amino acid sequence of LptM^His^ using amber-codon site-directed mutagenesis of pLptM^His^. D*lptM* cells were co-transformed with both the pLptM^His^ derivative plasmids and pEVOL-pBpF (Table 2). Cells were cultured in minimal media until mid-exponential phase, supplemented with 1 mM pBpa and 200 μM IPTG for 1.5h. Two identical cell culture aliquots were withdrawn, and one was kept on ice protected from light, whereas the other was subjected to UV irradiation (Tritan 365 MHB, Spectroline) for 10 min on ice.

Envelope fractions were prepared from both non-treated and UV irradiated cells and solubilized in 20 mM Tris-HCl, pH 8, 12% (w/v) glycerol, 4% (w/v) SDS, 15 mM EDTA and 2mM PMSF. After removing non-solubilized material by centrifugation, the supernatants were diluted in RIPA buffer (50 mM Tris/HCl pH 8, 150mM NaCl, 1% [v/v] NP-40, 0.5% [w/v] sodium deoxycholate, 0.1% [w/v] SDS) and subjected to Ni-affinity purification of LptM^His^ as previously described (Ranava et al., 2021).

### Molecular dynamics simulations

The top ranking AlphaFold2 models for LptDE and LptDE-LptM were used to seed MD simulations. The models were built into simulation systems using CHARMM-GUI (Jo et al., 2007; Lee et al., 2016a). Protein atoms were described with the CHARMM36m force field (Best et al., 2012; Huang et al., 2017). The C-terminus of LptE was truncated after Thr174 to remove contacts with the LptD barrel, and because the lDDT was low for this region. The N-terminal cysteine residues of LptM and LptE were tri-palmitoylated. Side chain pKas were assessed using propKa3.1 (Søndergaard et al., 2011), and side chain side charge states were set to their default, apart from Glu263 and Asp266 of LptD, and Lys103 of LptE, which were all set to neutral. The proteins were built into asymmetric membranes, comprising 6:3:1 POPE, POPG, and cardiolipin in the inner leaflet, with LPS in the outer leaflet. The *E. coli* Lipid A core with two 3-deoxy-α-D-manno-octulosonic acid units was chosen as a representative LPS molecule. The membranes were solvated with TIP3P waters and neutralised with K^+^, Cl^-^ and Ca^2+^ to 150 mM. Each system was minimized and equilibrated according the standard CHARMM-GUI protocol. Production simulations were run in the NPT ensemble, with temperatures held at 303.5 K using a velocity-rescale thermostat and a coupling constant of 1 ps, and pressure maintained at 1 bar using a semiisotropic Parrinello-Rahman pressure coupling with a coupling constant of 5 ps (Bussi et al., 2007; Parrinello and Rahman, 1981). Short range van der Waals and electrostatics were cut-off at 1.2 nm. Simulations were run to 500 ns and in triplicate for each system.

All simulations were run in Gromacs 2020.1 (https://doi.org/10.5281/zenodo.7323409) (Abraham, 2015). Data were analyzed using Gromacs tools (including residue distances, H-bond number, RMSF and angle analysis) and visualized in VMD (Humphrey et al., 1996). Plots were made using Matplotlib https://ieeexplore.ieee.org/document/4160265.

## ACKNOWLEDGMENTS

Research in RI’s lab was funded by the CNRS (ATIP grant to RI), the China Scholarship Council fellowships to YY and HC, and the FRM postdoctoral fellowship to DR. We thank the LMGM, UMR5100, for access to the *E. coli* Keio collection. HDX-MS experiments were supported by the French Ministry of Research (Investissements d’Avenir Program, Proteomics French Infrastructure, ANR-10-INBS-08 to O. B-S) and the Région Midi Pyrénées to O.B-S. Research in PJS’s lab was funded by Wellcome (208361/Z/17/Z) and BBSRC (BB/P01948X/1, BB/R002517/1 and BB/S003339/1). This project made use of time on ARCHER2 and JADE2 granted via the UK High-End Computing Consortium for Biomolecular Simulation, HECBioSim (http://hecbiosim.ac.uk), supported by EPSRC (grant no. EP/R029407/1). This project also used Athena and Sulis at HPC Midlands+, which were funded by the EPSRC on grants EP/P020232/1 and EP/T022108/1. We thank the University of Warwick Scientific Computing Research Technology Platform for computational access.

## Notes

### Competing Interest Statement

The authors have declared no competing interest.

