## Supplemental Text for "LptM promotes oxidative maturation of the lipopolysaccharide translocon by substrate binding mimicry"

### SUPPLEMENTARY METHODS

#### Genome samples for taxonomic analysis

A representative set of 2927 bacterial genomes was assembled by selecting one genome per bacterial Family from the Genome Taxonomy Database (GTDB) (Parks et al., 2018) and downloaded from the NCBI website (ftp://ftp.ncbi.nlm.nih.gov/genomes). In GTDB, Beta-proteobacteria are classified as an order within the class Gamma-proteobacteria. Representative set of Alpha- and Gamma-proteobacteria down to the Genera level and *Enterobacteriaceae* samples down to the Genera and Species level were also built.

#### Search of LptM-like candidates

Genomes were annotated with *Prokka* in fast mode with default settings (Seemann, 2014). Annotation of proteins with Pfam domains (version 34.0) was performed with *hmmscan* (*HMMER* package version 3.1b2, (Eddy, 2011)) and results were filtered to keep only the best non-overlapping alignments. A first analysis showed that genes coding for LptM-like proteins were not always annotated by *Prokka*. Identified candidate sequences were biased in their amino acid composition: the N-terminal regions were enriched in hydrophobic amino acids and the C-terminal regions were reach in amino acid typical of intrinsically disordered protein segments.

Genes located at the ends of contigs may have partial sequences. To obviate these problems, we translated the genomes in all six phases with an ORF size greater than or equal to 10 nucleotides to maximize the probability of identifying candidate genes (*esl-translate* program from *HMMER* package). ORFs internal to *Prokka* annotated genes were excluded from the analysis.

Initially, the presence of the PF13627 domain was used to identify LptM protein candidates in the translated ORFs (trORFs) obtained using *hmmsearch* (from *HMMER* package -E 1). The results indicated that *lptM* gene candidates are generally present in a single copy per genome. By selecting only a sequences with the lowest E-value for each genome, the sample size was reduced to 985 hits, of which 292 are present in the 476 proteobacterial genomes and 693 in the 2451 non-proteobacterial genomes.

In a second step, *hmmscan* (from *HMMER* package) was used with the Pfam library (version 34.0) against the 985 trORFs selected with the PF13627 profile. 911 trORFs have PF13627 as the best profile which eliminates 74 sequences. The latter sequences are more frequent in non-proteobacterial genomes (9.8% *vs.* 2%) and are mainly annotated as lipoproteins. We estimated the length of the C-terminal region (defined as the amino acid sequence of protein downstream of the PF13627 domain) by subtracting the final position of the alignment with the PF13627 domain from the sequence size. The vast majority of candidate proteins annotated in Alpha- and Gamma-proteobacteria have short C-terminal region (<70 amino acids), whereas non-proteobacterial candidates present a longer C-terminal region (Figure S2A). We filtered out candidates with a C-terminal region length greater than or equal to 70 amino acids. In Figure S2B are plotted the distribution of scores and alignment lengths for the three groups of genomes.

Genomic context of proteobacterial *lptM* gene candidates was extracted from GFF3 *Prokka* annotation files over a 5000 nucleotides window upstream and downstream of these genes. A classification of the proteins encoded by the neighboring genes into groups of homologous sequence was performed with *mmseq2* (Hauser et al., 2016), followed by a partition of the graph in communities with the Leiden method (Traag et al., 2019) of the *igraph* package (https://igraph.org/r/). The genes of two most frequent protein clusters are located downstream and upstream the *lptM*-like genes. The average gap between gene clusters and candidate genes is less than 32 nucleotides, suggesting that in the majority of genomes these genes are part of the same operon. The first cluster belongs to the the Orn/Lys/Arg decarboxylase class-II family as suggested by the presence of the domain PF02784. The second cluster include genes encoding Lyases (PF00206). This conservation of neighboring gene is more common in Gamma- than Alpha-proteobacteria (Figure S2C). A phylogenetic tree was calculated on the 2927 genomes of our bacterial sample (Figure S2D). The alignment of the 120 markers of these genomes was retrieved from the GTDB.

The tree was inferred with *fasttree* (Price et al., 2009). Branch supports were estimated with the local bootstrapping. The logoplots (Figure S2E) of the LptM-like proteins identified in Gamma-, Alpha-, and non-proteobacteria were calculated with *R* package *ggseqlogo* (Wagih, 2017).

To obtain a more accurate picture of the distribution and evolution of *lptM* in *Enterobacteriaceae*, we extended our analysis to 766 *Enterobacteriaceae* species. We identified 545 LptM-like proteins. To characterize the C-terminal regions of LptM, we used the *meme* software (-protein -mod zoops -nmotifs 25 -minw 4 -maxw 16 -minsites 3 -evt 0.05), which identifies blocks of conserved motifs in a subset of sequences (Bailey et al., 2009). To obtain a better taxonomic representation, we selected one sequence from each genus (93 sequences). The motifs detected by *meme* were annotated on the whole sequences with the *mast* software (Bailey et al., 2009). The maximum motif size of 16 AA covers the lipobox and the downstream conserved region.

#### Analysis of LptM conservation in *Enterobacteriaceae*

In Alpha- and Gamma-proteobacteria, two genes are frequently conserved upstream and downstream of the *lptM*-like genes (Figure S2C). This conservation of a chromosomal neighboring genes surrounding *lptM* candidates suggests that they were inherited from a common ancestor and are therefore orthologs. This reinforces the hypothesis that, in proteobacteria, the PF13627 domain reliably identifies proteins that have a common origin and similar functions. This also suggests that the proteins predicted from the translated ORFs are functional.

The distribution of LptM-like in bacteria shows that they are very frequent and with high scores in Gamma-proteobacteria (in 82.1% of genomes), less frequent with lower scores in Alpha-proteobacteria (61.4%) and infrequent (9.1%) with very low scores in non-proteobacteria (Figure S2D). By filtering the results with a thresholds score ≥ 13 and an alignment length score ≥ 16, the number of candidates in non-proteobacteria and in Alpha-proteobacteria is significantly reduced whereas the number of candidates in Gamma-proteobacteria is marginally reduced (1.7%, 56.6% and 78.9%, respectively). In agreement with these observations, LptM candidates from Gamma-proteobacteria show more extensive sequence conservation downstream of the the lipobox cysteine (Figure S2E). In addition, *lptM*-like genes appear to be randomly distributed in non-proteobacterial genomes. This distribution and the low sequence scores suggest that they do not encode for a true homolog of proteobacterial LptM. However, these non-proteobacterial proteins have lipoprotein characteristics captured by the PF13627 profile.

We identified the LptD, LptE, and LptM proteins in the genomes of 776 species of *Enterobacteriaceae*. The results are summarized in Figure S3, where only the 135 representative genomes of the GTDB *Enterobacteriaceae* genera are shown. Based on phylogenetic analyses and the use of conserved molecular features it was proposed to divide the *Enterobacteriaceae* family into seven new families (Adeolu et al., 2016): a restricted *Enterobacteriaceae* family, and the new families Erwiniaceae, Pectobacteriaceae, Yersiniaceae, Hafniaceae, Morganellaceae and Budviciaceae. The newly defined family of restricted *Enterobacteriaceae* was subdivided into 6 subfamilies: *Escherichia*, *Klebsiella*, *Enterobacter*, *Kosakonia*, *Cronobacter*, *Cedecea*, and an "*Enterobacteriaceae incertae sedis*" clade containing species whose taxonomic placement within the family is not clear (Alnajar and Gupta, 2017). The phylogenetic tree reconstructed on the 135 representative genomes is in agreement with this classification (Figure S3).

As above, LptM proteins are not always annotated as genes but can be identified as translation products of ORFs. We annotated LptD and LptE proteins in these genomes using PF04453 and PF04390 profiles. The three proteins are co-occurring in the vast majority of genomes larger than $2x10^6$ nucleotides, while they are present in only 6 of 33 genomes smaller than $2x10^6$ nucleotides. In these genomes LptM is most often absent and LptD is most often present. The bacteria that underwent a strong reduction of their genome size belong to subtrees that include endosymbiotic bacteria (Figure S3). The reduction in genome size is accompanied by the loss of a large number of genes, so the simultaneous loss of the genes encoding the three Lpt proteins in 14 out of 33 genomes provides only weak evidence for the existence of a functional link between these three proteins.

Two *meme* motifs are found exclusively in the LptM proteins of genomes belonging to the newly defined restricted *Enterobacteriaceae* family (Figure S3). This distribution suggests that they were acquired in the last common ancestor of this subfamily, either in a single event or stepwise. The presence of these motifs stabilized the C-terminal region which is generally variable in length and sequence in other LptM proteins. The third motif which is rich in polar amino acids (Q, N, T and S) is present in proteins of other families with a low frequency (18/70).

Outside the *Enterobacteriaceae* family, LptM proteins lack sequence conservation for the C-terminal region, suggesting that either this segment of LptM does not mediate specific protein-protein interactions in these bacteria or that the nature of these interactions is not evolutionarily conserved. Instead, the sequences of LptM in the restricted *Enterobacteriaceae* family presents several distinguishing motifs including the presence of a conserved C-terminal region, suggesting a relatively recent acquisition.

### SUPPLEMENTARY FIGURE LEGENDS

**Figure S1. MALDI-TOF mass-spectrometry analysis of purified LptDE$^{His}$.**

Gel bands obtained with BN gel electrophoresis of LptDE$^{His}$ complex purified from wild-type (top spectrum) or Δ*lptM* (bottom spectrum) cells were subjected to in-gel trypsin digestion and MALDI-TOF analyses. LptD and LptE were identified by peptide mass fingerprinting. The m/z values of peaks matching LptD or LptE peptides are indicated in blue and brown respectively (top and bottom spectra). Comparison of the two MS spectra led to the identification of discriminant peaks between the wild-type and the Δ*lptM* samples (m/z values in red). MALDI TOF/TOF fragmentation of ion parent at m/z= 2520.28 (indicated by a red arrow) allowed the identification of the peptide $_{34}$NAPPPTKPVETQTQSTVPDKNDR$_{56}$ of LptM. y and b ions resulting from fragmentation are indicated (central spectrum). Internal fragment ions are labeled with one or two asterisks.

**Figure S2: Distribution of LptM-like proteins in bacteria.**

LptM-like proteins were predicted in the translated ORFs of represented families of bacteria with *hmmsearch* and the PF13627 domain.

**(A)** Length distribution of the C-terminal region of the sequences for non-proteobacteria and Alpha- and Gamma-proteobacteria (vertical bar at 70 AA).

**(B)** Alignment score and length distribution in the three samples. Vertical bars indicate strict score (13) and length (16) thresholds.

**(C)** Venn diagram with the occurrence of PF00206 and PF02784 protein-coding genes in the neighborhood of *lptM*-like genes in Alpha and Gamma-proteobacteria.

**(D)** Inferred phylogenetic tree for representative bacterial families with *fasttree* from the alignment of 120 markers retrieved from GTDB. The color gradient of the branches is proportional to the local bootstrap support values from 0 (red) to 1 (purple). Alpha and Gamma-proteobacteria clades are indicated on the tree (yellow and blue shading of subtrees). To simplify the annotation, only phyla that contain at least 10 genomes have been indicated in the figure on the first ring with a color code and their names are reported in the outer ring. Genome size is shown as a purple histogram with a threshold at $2x10^6$ nucleotides, i.e., genomes smaller than $2x10^6$ nucleotides are shown as negative. The next ring shows the value of the LptM-like protein score with a threshold of 13 (scores below 13 are shown as negative).

**(E)** Sequence logos of LptM-like proteins identified in Gamma-, Alpha- and non-proteobacteria.

**Figure S3: Distribution of LptD, LptE and LptM proteins in *Enterobacteriaceae* and conserved motifs in LptM.**

We used a single representative genome per genus of *Enterobacteriaceae*. Left column, the tree calculated on the alignment of the 120 markers retrieved from GTDB with IQ-TREE (Minh et al. 2020). The lengths of the branches are not proportional to the distances inferred by IQ-TREE, because genomes (including endosymbiont) with very high evolutionary rates mask the other parts of the tree. The branch supports were assessed with the SH-like approximate likelihood ratio test (-alrt 1000). It was reported on the tree as branch color gradient. The tree is rooted using the tree obtained with the representative genomes of the Bacteria families (Figure S2D). The division of the GTDB family *Enterobacteriaceae* into seven families (Adeolu et al., 2016) and the subdivision into six subfamilies of the newly defined family of restricted *Enterobacteriaceae* (Alnajar and Gupta, 2017) were reported on the tree. Second column, histogram of the genome sizes. A threshold of $2x10^6$ nucleotides was used to highlight the presence of small genomes. Third column, presence/absence of LptD, LptE and LptM proteins. In the LptM column the dark red circles indicate the presence of an annotated gene and the lighter circles an ORF whose translation product is similar to LptM. Last column, LptM mast annotation of the four identified meme domains. The sequence logos of the motifs are indicated. The *Escherichia* genus, represented by *Escherichia fergusoniiis*, is located at the bottom of the tree (colored in red).

The terms endosymbiont, endobia and symbiotica present in the genome descriptions are colored in blue.

**Figure S4**. **Site specific photo-crosslinking of LptM.**

**(A)** Δ*lptM* cells harboring pEVOL-pBpF and expressing $LptM^{His}$ derivative forms with pBpa engineered at specific amino acid positions were subjected to UV irradiation as indicated and envelope fractionation (Load) followed by $LptM^{His}$ affinity purification (Elution). Samples were analyzed by SDS-PAGE and Western blotting using the indicated antisera. Load: 2.2%; Elution: 100%.

**(B)** Upon UV irradiation, the envelope (Load) and the elution fractions obtained from Δ*lptM* cells expressing $LptM^{His}$ containing pBpa at position L22 was analyzed by SDS-PAGE and Western blotting using an anti-LptM antiserum. Load: 2.2%; Elution: 100%.

**(C)** Upon UV irradiation, as indicated, $LptM^{His}$ containing pBpa at position V42 was purified from Δ*lptM* or Δ*lptM ompA::kan* cells and analyzed by SDS-PAGE and Western blotting using anti-His and anti-OmpA antisera.

**Figure S5**. **Structure prediction and molecular dynamics simulations of LptDE and LptDEM.**

**(A)** A view of the LptD hinge in the LptDE-LptM AphaFold2 model. The N-terminal Cys position is shown, as are the predicted positions of LptD C1 to C4. Graphs plot the minimum distance between selected Cys pairs throughout 3 x 500 ns simulations of the LptDE-LptM system (each run plotted separately in red, blue or orange).

**(B)** Angle of LptD β-taco in relation to the LptD β-barrel, as computed using vectors between the residues shown in cyan. The presence of LptM shifts the angles to be lower (more upright).

**(C)** Top: C-alpha RMSFs over 3 x 500 ns simulation projected onto the LptDE or LptDE-LptM model. Bottom: RMSF plots focused on the LptD N-terminus, which is destabilized by LptM.

**(D)** A zoom-in of the LptDEM heterotrimer modeled using AlphaFold2 (as in Figure 5A) highlights the interaction of a LptM N-terminal segment with the LptD β-barrel domain. A salt-bridge is predicted to form between LptM K23 and LptD E275.

**Figure S6**. **LptM interacts with LptE and LptD β-barrel domain.**

The LPS translocon was purified by Ni-affinity purification of $LptM^{His}$ from Δ*lptM* cells transformed with $pLptDEM^{His}$ or $pLptD^{\beta\text{-barrel}}EM^{His}$, as indicated.

**Figure S7**. **HDX-MS of the LptDE translocon.**

**(A)** Sequence coverage maps of LptD (top) and LptE (bottom) obtained by HDX-MS of LptD and LptE. The color scale shows the relative deuterium uptake after 30 sec of deuteration from 0% (blue) to 60% (red).

**(B)** Relative deuterium uptake of LptDE after 30 sec deuteration color coded from 0% (blue) to 60% (red) on the three-dimensional structure of LptDE, as predicted by our AlphaFold2 model.

**Figure S8. Deuteration heatmap of LptD and LptE**

**(A)** Differential heatmap of LptD (top) and LptE (bottom) between the LptDE translocon in the presence or absence of LptM color coded from -35% (red: protected in the complex)/0% (white: no difference)/+35% (blue: deprotected in the complex).

**(B)** The color coded heatmap described in (A) is represented on the three-dimensional structure of LptDE, as predicted by our AlphaFold2 model.

**Figure S9. LptDE heterodimer and LptDEM heterotrimer deuteration Wood Plots.**

Differential Woods Plots between LptDE heterodimer and LptDEM heterotrimer, at the different time points of deuteration (0.5, 1, 5, 10 and 30min), showing in blue and red, peptides that are significantly protected and deprotected, respectively (hybrid significance test, p-value<0,001, Confidence Interval 0,52 Da).

**Figure S10. Mass spectra of LptD β-taco peptides showing EX1 kinetics.**

For each peptide are shown: top, Relative deuterium uptake as a function of deuterium exposure time (red line for the LptDE and blue line for the LptDEM samples); bottom: mass spectra obtained with LptDE (left) and LptDEM (right).
