## Supplemental Figures for "LptM promotes oxidative maturation of the lipopolysaccharide translocon by substrate binding mimicry"

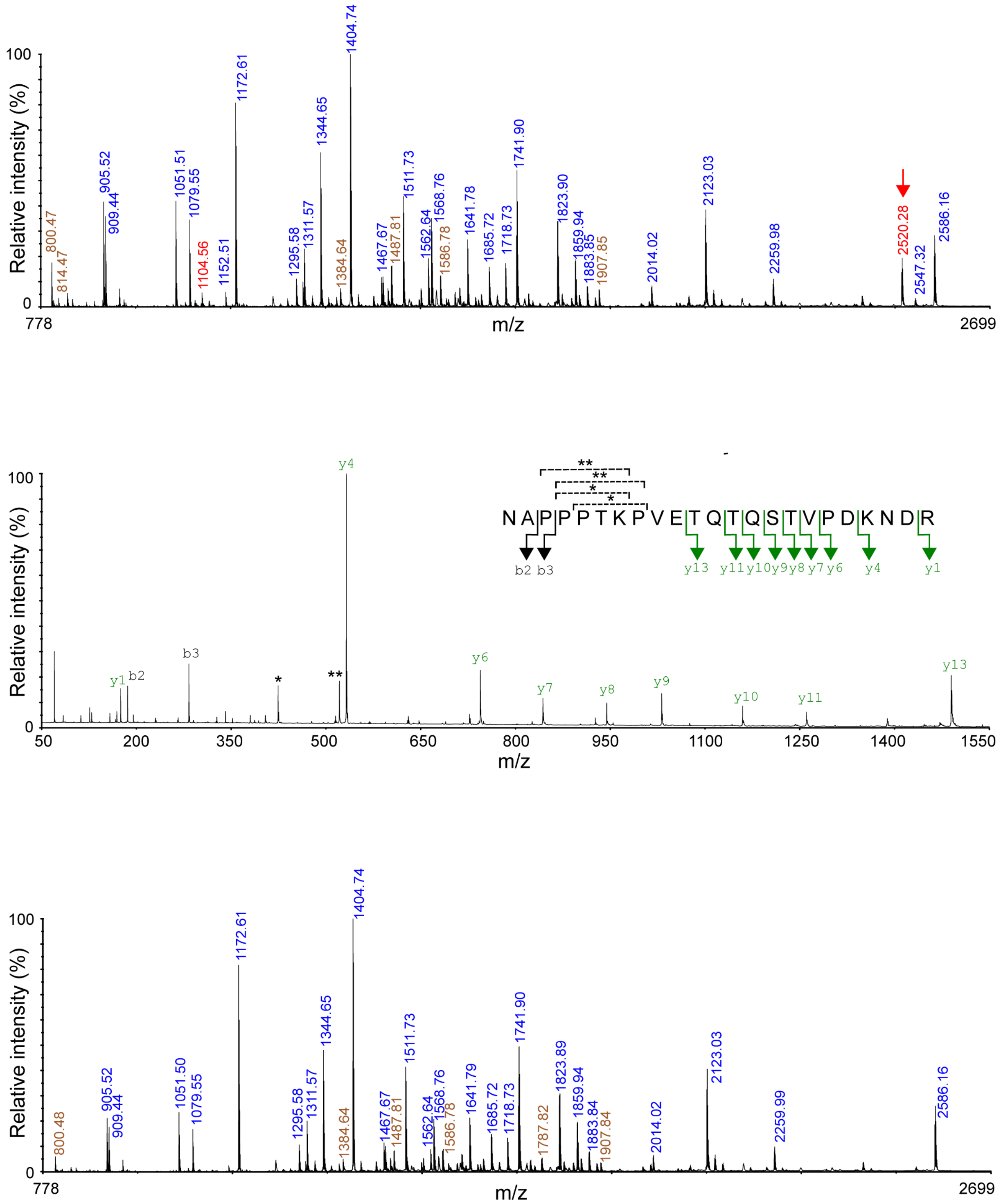


Yangt et al., Figure S1

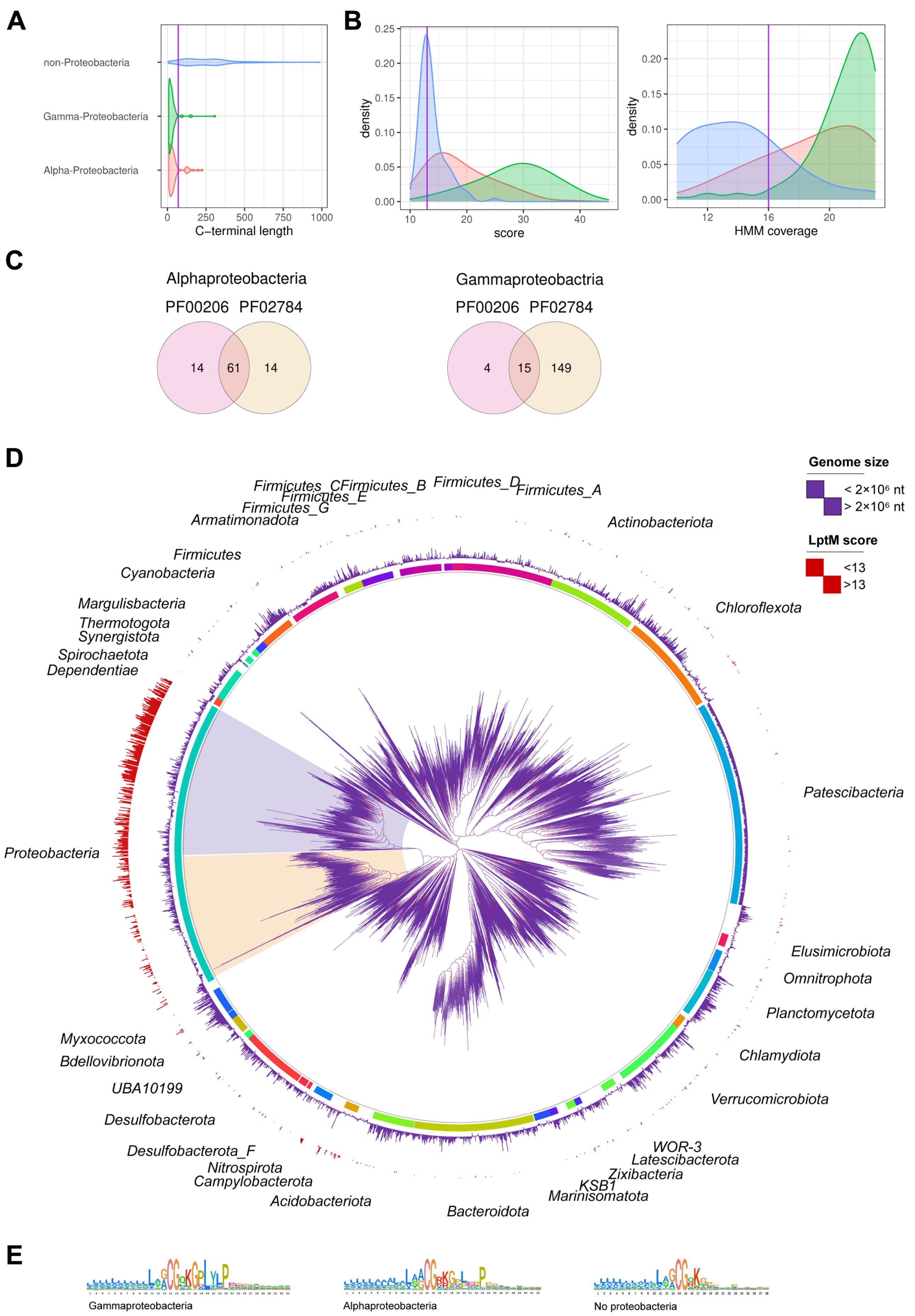


Yang et al., Figure S2

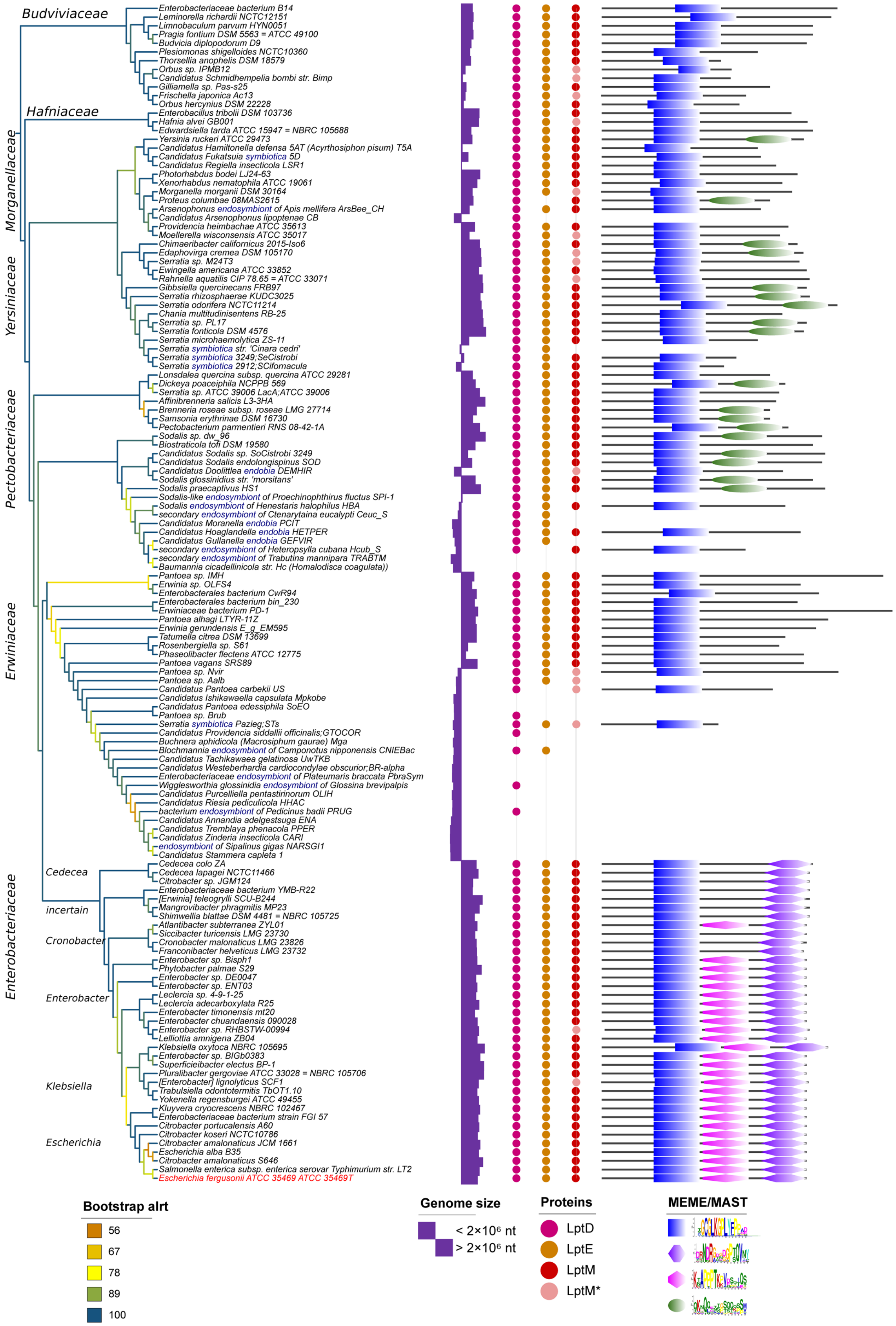


Yang et al., Figure S3

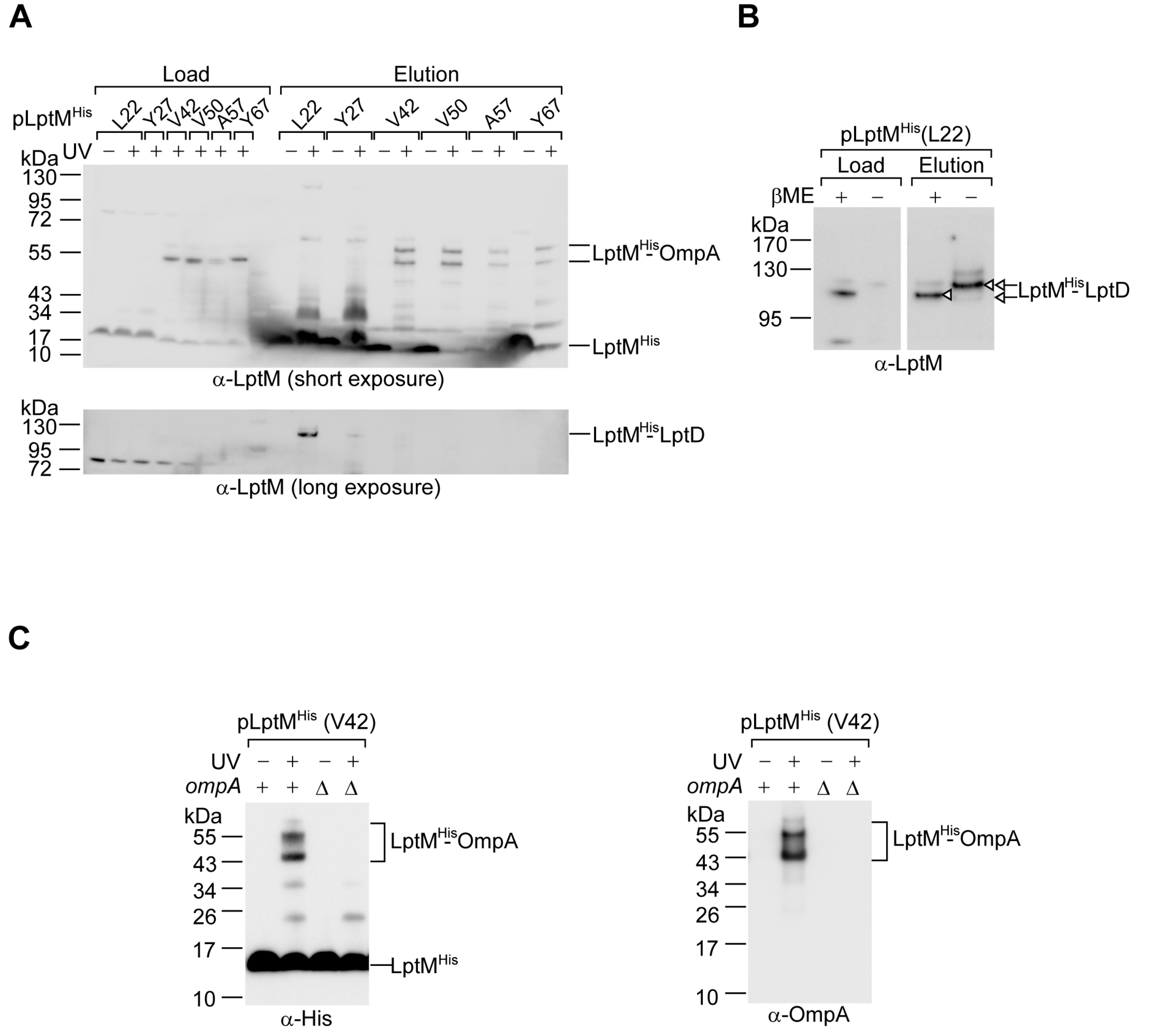


Yang et al., Figure S4

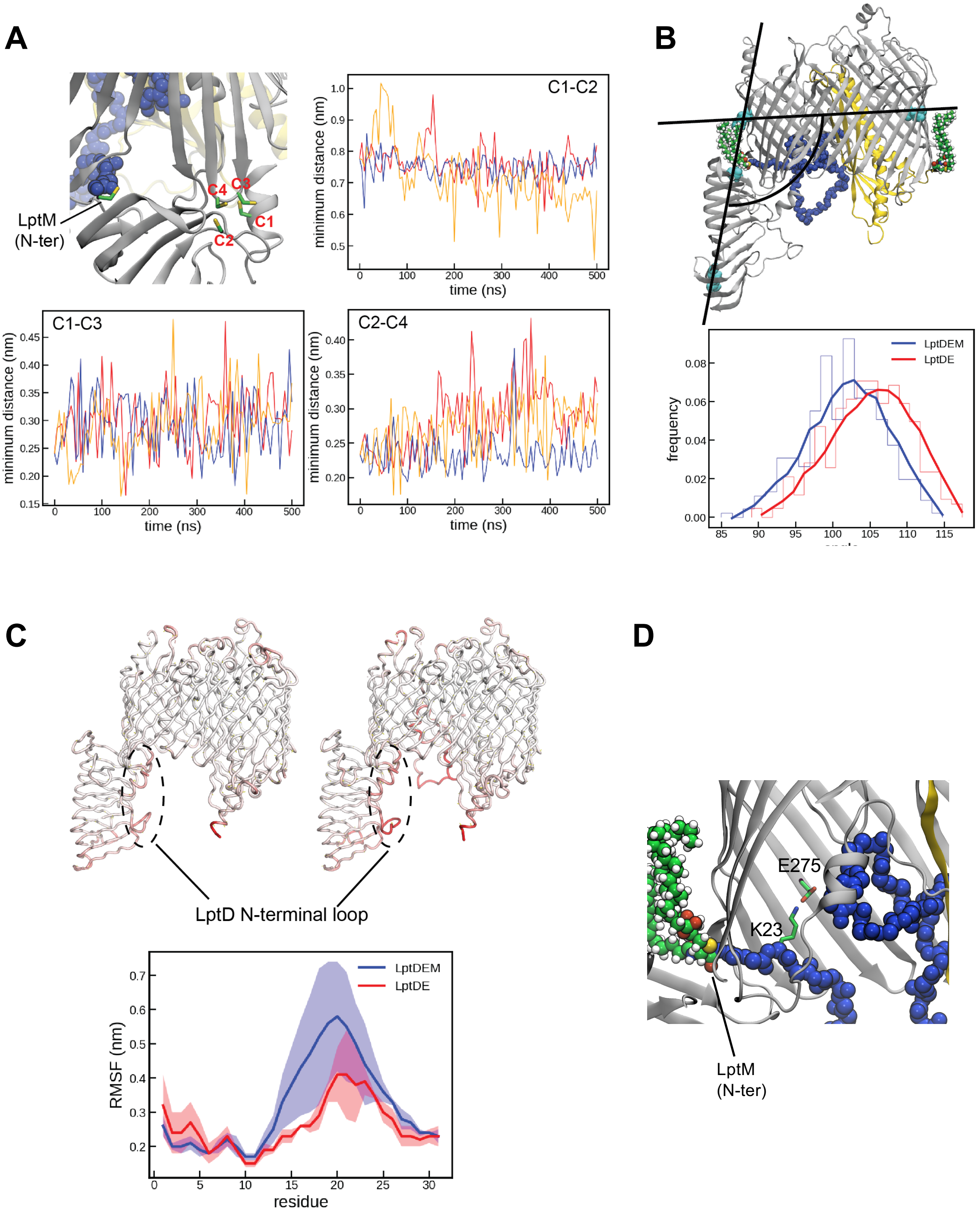


Yang et al., Figure S5

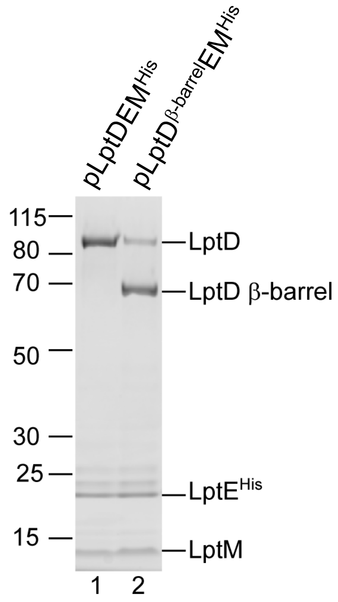


Yang et al., Figure S6

A

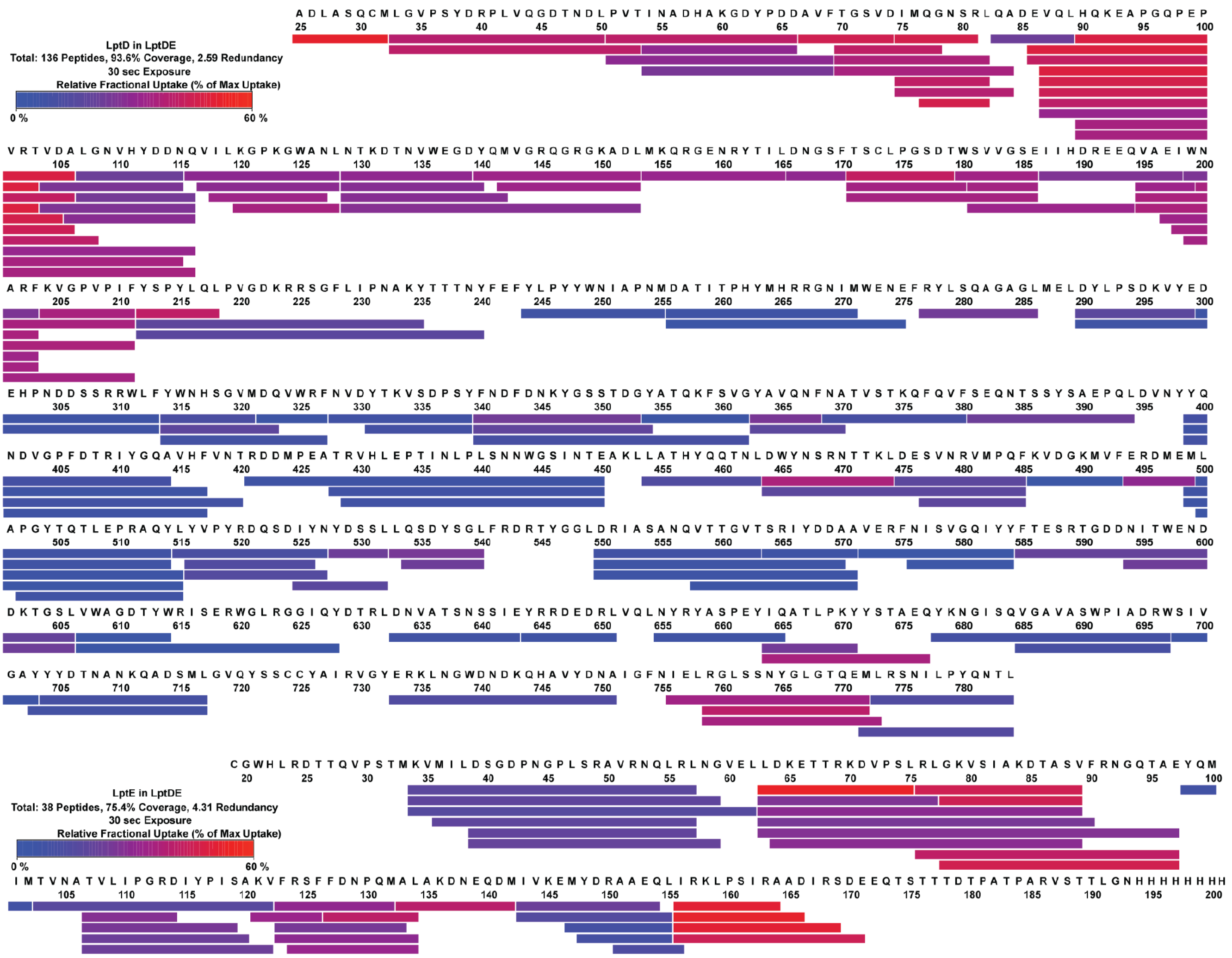


B

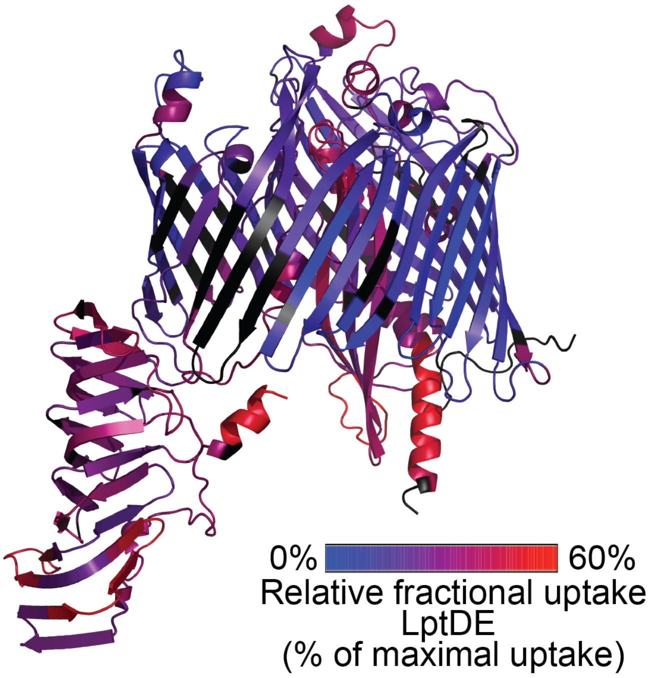


Yang et al., Figure S7

**A**

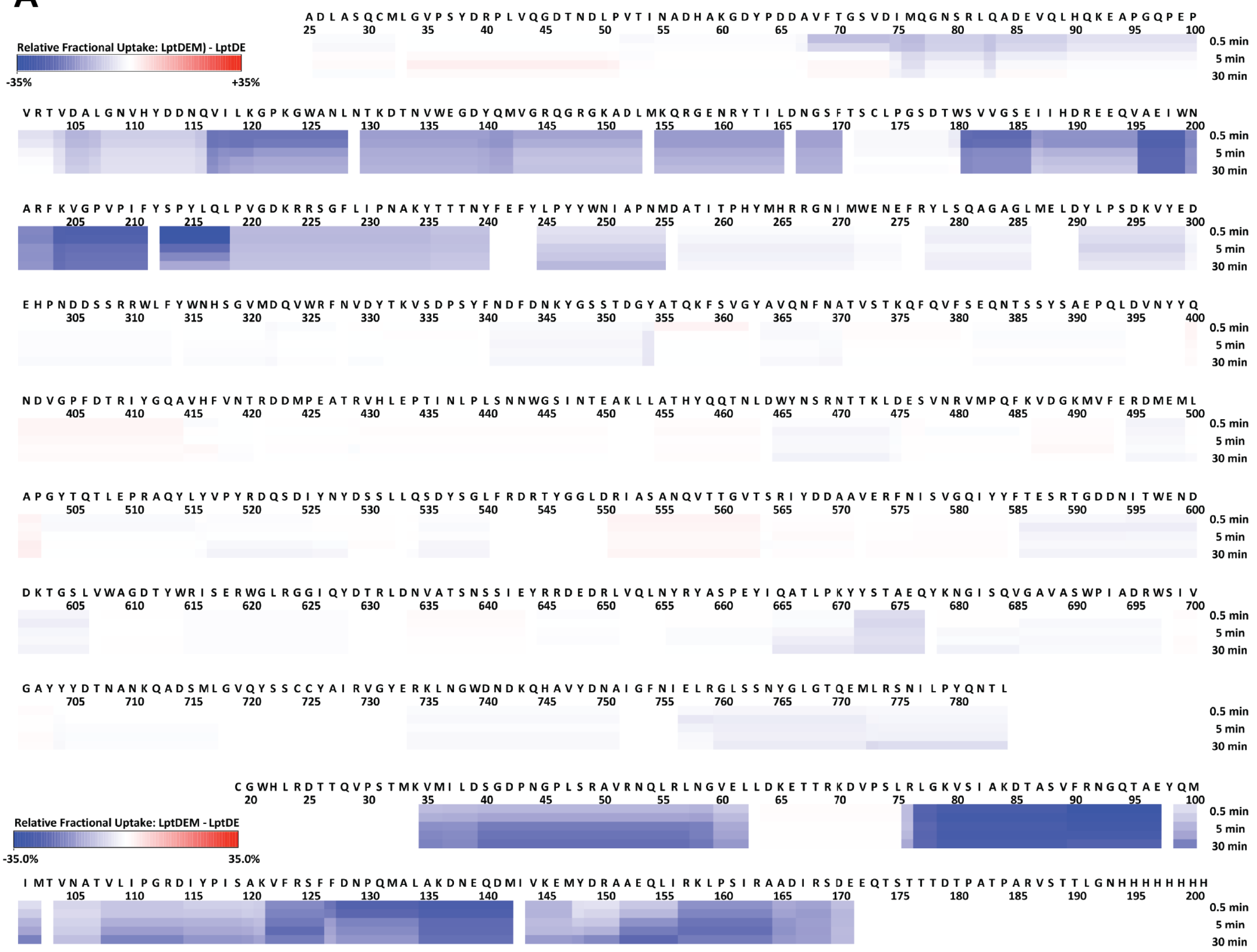


**B**

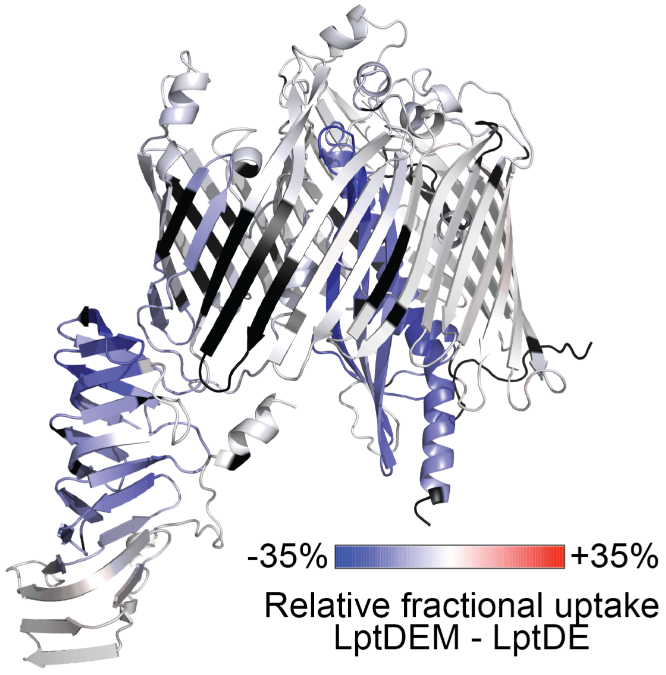


Yang et al., Figure S8

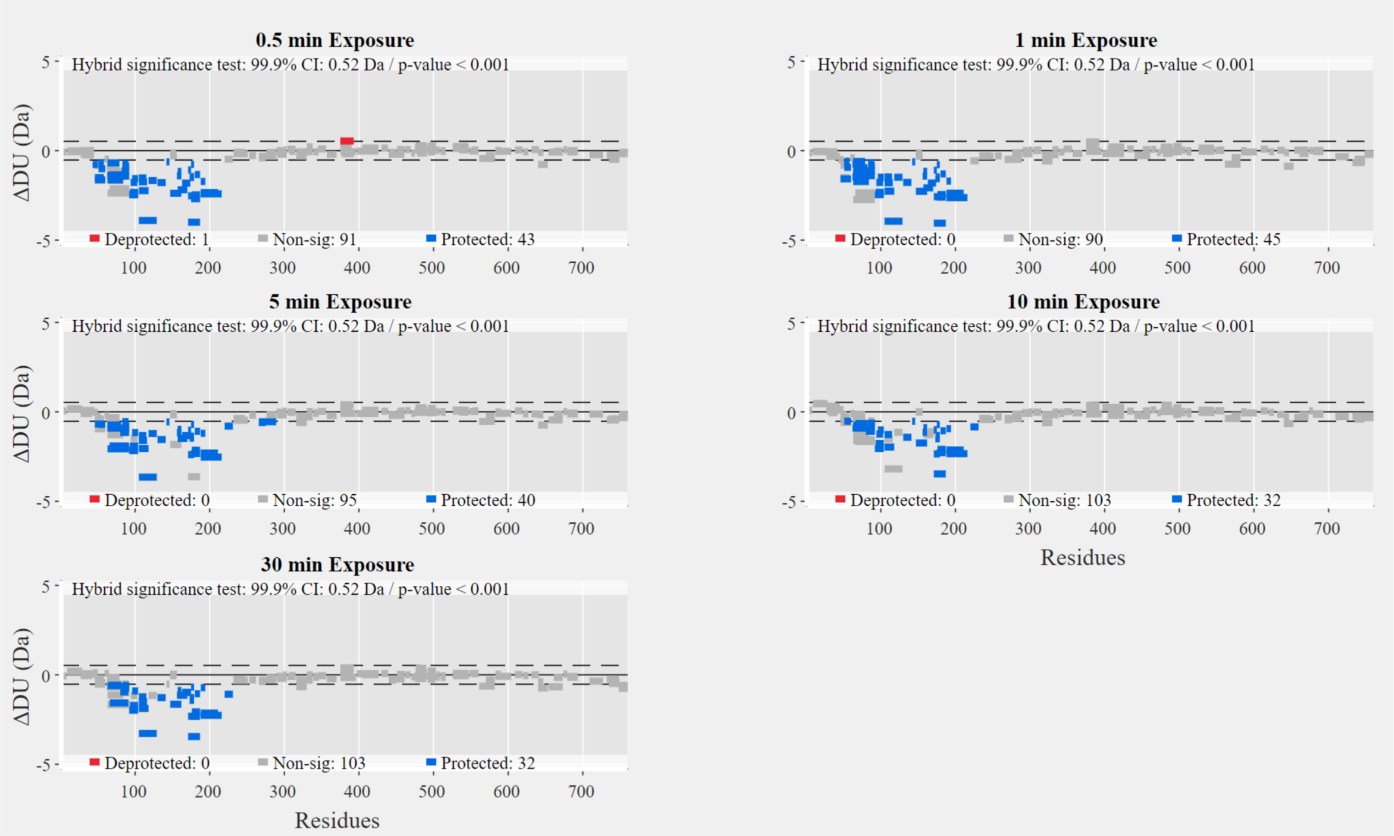


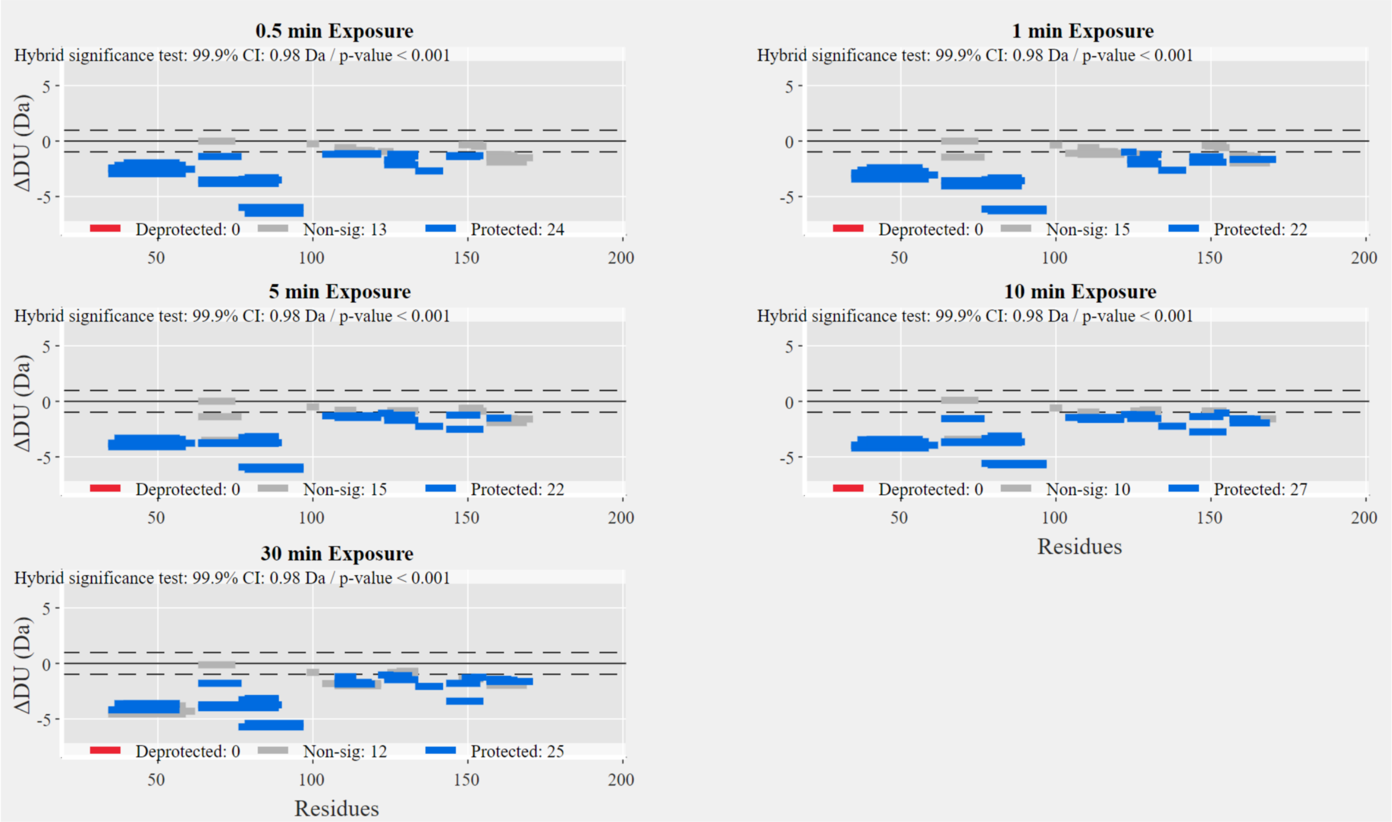


Yang et al., Figure S9

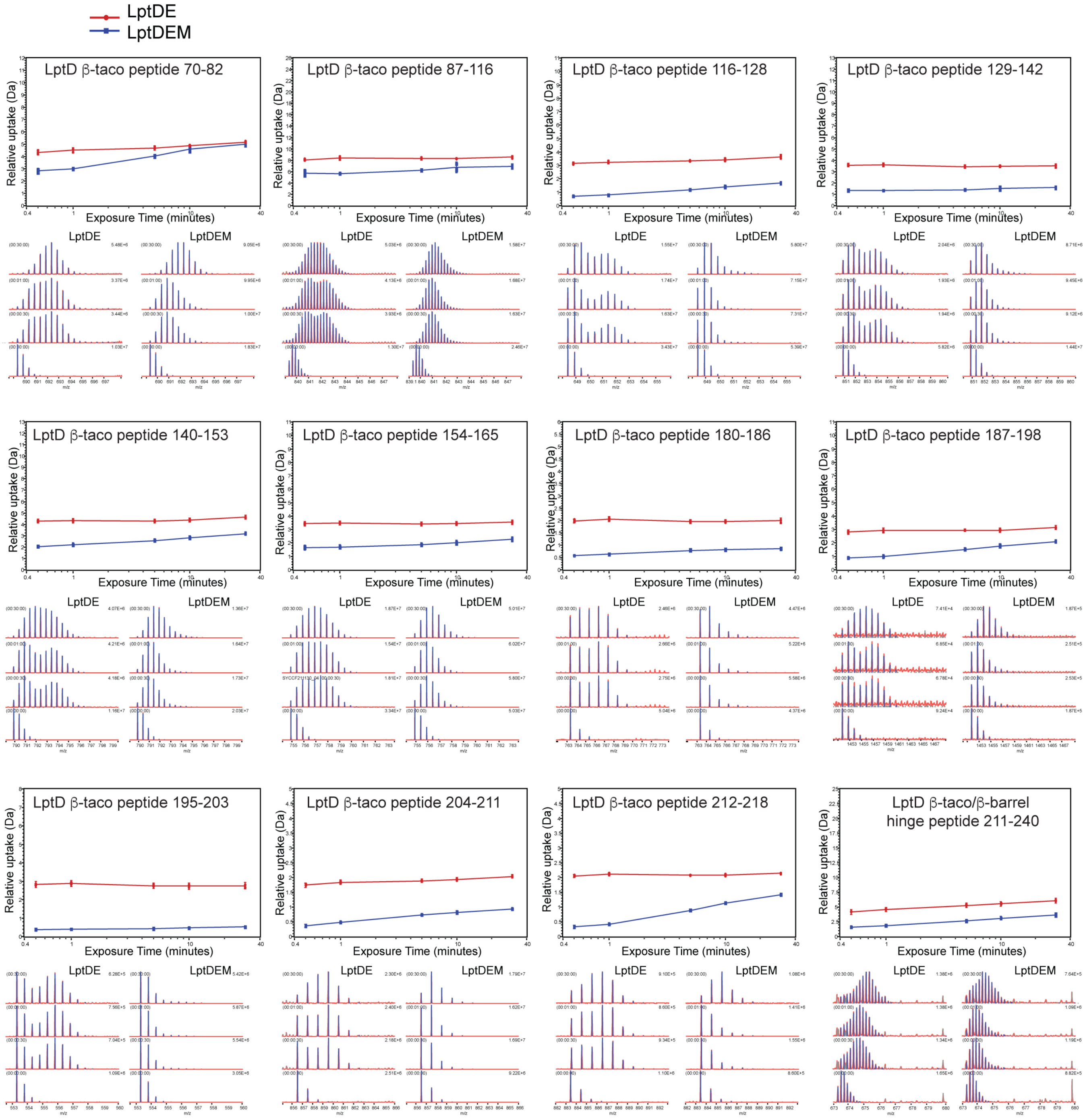


Yang et al., Figure S10
